# Interprotomer Communication and Functional Asymmetry in H/ACA snoRNPs

**DOI:** 10.1101/2025.06.07.658439

**Authors:** Hemendra Singh Panwar, Timothy John Vos, Xiaoyan Xie, Hyo Sik Jang, Hyoungjoo Lee, Ryan Sheldon, Evan Worden, Ute Kothe

## Abstract

H/ACA small nucleolar ribonucleoproteins (H/ACA snoRNPs) facilitate essential cellular processes such as RNA modification, folding, and stability. Here, we present multiple cryo-EM structures of endogenous, catalytically active insect H/ACA snoRNPs containing two protomers assembled on a two-hairpin H/ACA snoRNA. By characterizing key protein-protein and protein-RNA interactions, we reveal the coordination of pseudouridylation activity across the two protomers which explains the predominance of two-hairpin structures in eukaryotic H/ACA snoRNAs. Moreover, we discovered that several mutations in H/ACA proteins associated with Dyskeratosis congenita directly impair pseudouridine formation suggesting how these mutations disrupt RNA modification and ribosome biogenesis in this disease. Additionally, we uncover coordinated structural changes between Nop10, Nhp2 and the N-terminal extensions of Cbf5 in the 3′ protomer that resemble active and inactive conformations and may regulate H/ACA snoRNP activity. In summary, this study provides detailed insight into the structure and function of RNA modification-competent, asymmetric H/ACA snoRNPs, which play pivotal roles in cellular processes including ribosome biogenesis, rRNA folding, (m)RNA modification, and telomere maintenance.

**Significance statement:** H/ACA snoRNPs are critical for numerous RNA processes as they utilize different H/ACA snoRNAs to introduce pseudouridines in ribosomal and other non-coding RNAs as well as mRNAs. In addition, H/ACA snoRNPs stabilize and fold RNA thereby contributing to ribosome biogenesis and telomere maintenance. Mutations in H/ACA snoRNP subunits such as dyskerin (pseudouridine synthase), Nhp2, and Nop10 cause the genetic disorder Dyskeratosis congenita. Here, we present cryo-EM structures of active eukaryotic H/ACA snoRNPs, revealing an asymmetric, dimeric complex of two protomers. We identify the key inter-protomer contact hotspots and biochemically characterize mutations at these interfaces. Our findings emphasize the importance of inter-subunit interactions for snoRNP stability and activity, providing mechanistic insight into H/ACA snoRNP function and Dyskeratosis congenita pathogenesis.

## Introduction

H/ACA small nucleolar ribonucleoproteins (snoRNPs) are versatile molecular machines responsible for the site-specific isomerization of uridine to pseudouridine (Ψ) at many sites in ribosomal RNA (rRNA), small nuclear RNAs (snRNAs) and other RNAs (1). For this reason, H/ACA snoRNPs are critical for ribosome biogenesis in eukaryotes. In addition, specific H/ACA snoRNAs mediate functions unrelated to pseudouridine formation. For example, U17/snR30 is essential for biogenesis of the small subunit of the ribosome by contributing to the correct folding of 18S rRNA (2–4). Furthermore, the vertebrate telomerase RNA (hTR) contains a 3′ H/ACA domain, which is important for stabilizing and localizing telomerase (5–7). H/ACA snoRNAs are involved in other cellular processes, e.g. in protein secretion (8), and additional functions may be discovered as interacting RNAs remain unknown for some so-called orphan H/ACA snoRNAs (9). The overall cellular importance of H/ACA snoRNPs is also reflected in their association with human diseases (1).

In eukaryotes, the H/ACA snoRNA almost always has a conserved hairpin-hinge-hairpin structure where both hairpins direct pseudouridylation in an RNA-dependent, sequence-specific manner. The uridine to be modified is selected by the sequences flanking the uridine on either side, which base pair with a so-called pseudouridylation pocket in H/ACA snoRNA, while the central NU dinucleotide including the target uridine is left unpaired (**Fig. 1a**) (10). Immediately 3′ of each hairpin in H/ACA snoRNAs there is a conserved ANANNA (H box; 5′ hairpin) or ACA motifs (3′ hairpin) (**Fig. 1a**) which are the only characteristic sequence motifs found in all H/ACA snoRNAs (11). This two-hairpin structure is functionally important for eukaryotic H/ACA snoRNPs as truncation of the eukaryotic H/ACA snoRNA to a single hairpin reduces pseudouridylation activity (12).

**Figure 1.**
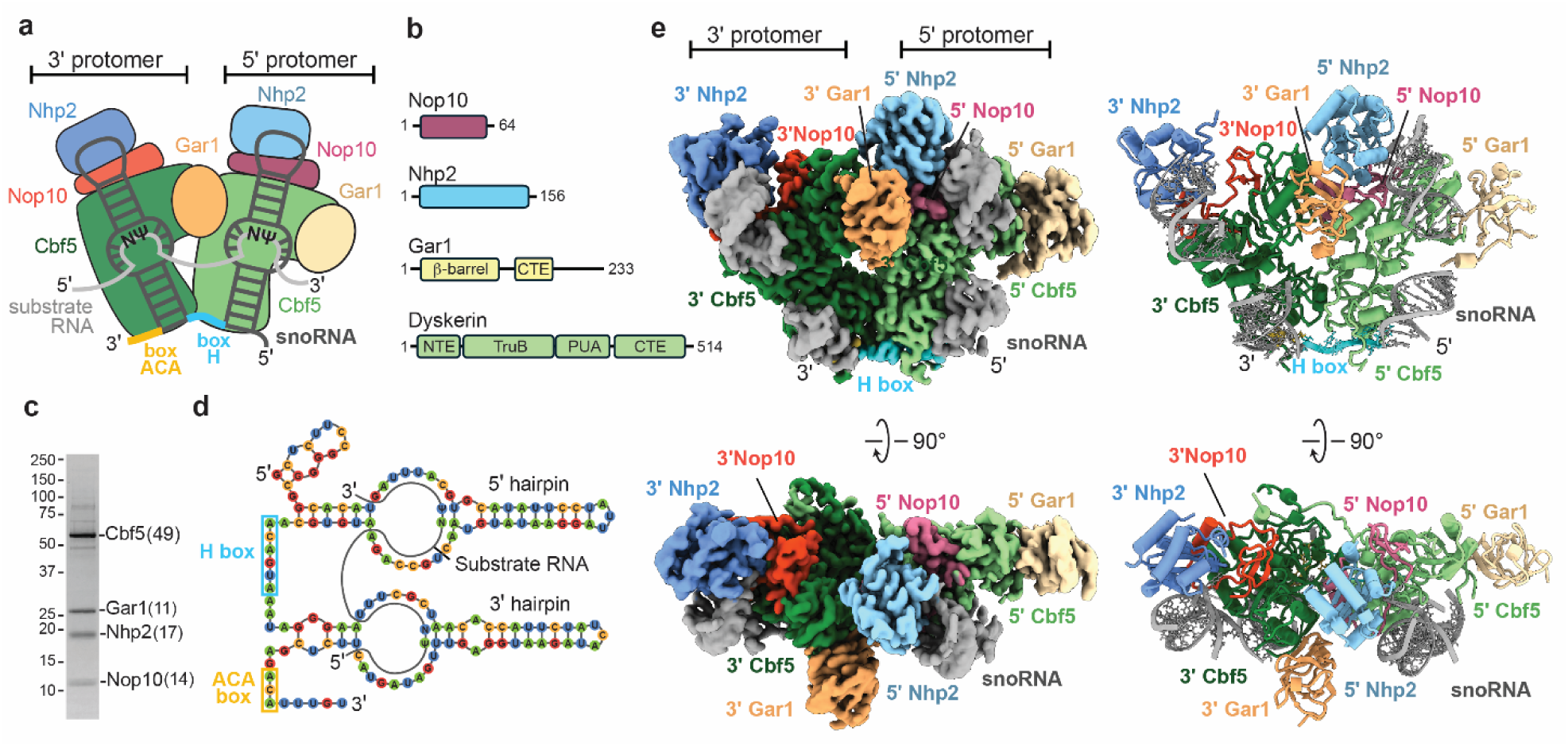
Overall architecture of Insect H/ACA snoRNPs. **(a)** Cartoon representation of the dimeric insect snoRNP complex - the four proteins in 5’ protomer are color indicated as - Cbf5 (light green), Gar1 (light yellow), Nop10 (dark magenta), and Nhp2 as (light blue) and in the 3’ protomer as Cbf5 (forest green), Gar1 (sandy brown), Nop10 (orange red), and Nhp2 (corn flour blue). The snoRNA (gray) is shown as a double stem loop structure bound to each protomer and connected by the H box (blue) and ending in an ACA box. Also, the substrate RNA (white) is shown bound to snoRNA. **(b)** Schematic representation of snoRNA proteins including their domains. **(c)** SDS-PAGE analysis of the purified insect H/ACA snoRNP complex. Number of peptides of each protein subunit identified by mass-spec is in parentheses. **(d)** Secondary structure of the most abundant snoRNA bound to substrate RNA - 2 stem loops (composed of stem and hairpin) with the conserved H box and ACA box is indicated. **(e)** 2.8 Å cryo-EM reconstruction (left) of the dimeric insect H/ACA snoRNP complex (class I) in different orientations. Also shown is the ribbon representation of the H/ACA snoRNP complex (right). The subunits are colored as indicated.

Previous structures of the single-hairpin archaeal H/ACA RNP have revealed the overall architecture of the complex and its interaction with RNA substrates (13, 14). Each hairpin of the H/ACA snoRNA binds to a protein complex comprising dyskerin (Cbf5 in yeast), Nop10, Gar1 and Nhp2 (L7Ae in archaea) (**Fig. 1a**). Cbf5/dyskerin provides the catalytic activity to H/ACA snoRNPs (15) and is composed of four domains: the pseudouridine synthase and achaeosine transglycosylase (PUA) domain, the tRNA pseudouridine synthase B-like (TruB) domain, the N-terminal extension (NTE), and a C-terminal extension (CTE) (**Fig. 1b**). The PUA domain of Cbf5/dyskerin binds to the ACA and H boxes (13, 16) and positions the catalytic domain of Cbf5/dyskerin next to the pseudouridylation pocket which targets a uridine in the substrate RNA for modification (12). Nop10 binds to the snoRNA and is sandwiched between Cbf5/dyskerin and Nhp2 (14, 16). Nhp2 interacts with the upper stem of the H/ACA snoRNA, and with Nop10, positions the H/ACA snoRNA on the catalytic Cbf5/dyskerin subunit (12, 17). Gar1 interacts with Cbf5/dyskerin without contacting the H/ACA snoRNA, but Gar1 indirectly mediates substrate RNA binding via its interactions with the Cbf5/dyskerin thumb loop (13, 14, 16, 18).

Cryo-EM structures of the human telomerase complex revealed the asymmetric assembly of the eukaryotic two-hairpin H/ACA snoRNP protomers (18–20). The 3′ hairpin protomer binds hTR through dyskerin, Nop10, and Nhp2 in a similar manner as the archaeal snoRNP (17, 19). However, the 5′ protomer interacts with hTR only through dyskerin.

Moreover, the two dyskerin proteins interact asymmetrically through their PUA domain. A similar protein arrangement on a two-hairpin H/ACA snoRNA has recently been reported for the yeast snR30 H/ACA snoRNA that is critical for rRNA folding (2). However, neither the telomerase nor the snR30 H/ACA snoRNP complex are active in pseudouridylation. Therefore, these structures do not address why eukaryotic H/ACA snoRNAs typically have two hairpins or how the two-hairpin structure might enhance H/ACA snoRNP pseudouridylation activity (12). In particular, it remains unclear how interactions across the two protomers affect catalysis of pseudouridine by the two dyskerin/Cbf5 proteins.

Mutations in dyskerin, Nop10, and Nhp2 give rise to an inherited disease called Dyskeratosis congenita characterized by bone marrow failure, pre-mature aging and resulting in childhood fatality in the most severe cases (21). In general, Dyskeratosis congenita is considered a telomerase disorder, but ribosome biogenesis and ribosome function are also affected in patients with mutations in dyskerin, Nop10, and Nhp2 (22, 23). Most Dyskeratosis congenita mutations occur within the dyskerin PUA domain and the N- or C-terminal extensions. However, a smaller subset of disease mutations are found within Nop10 and Nhp2 or the catalytic TruB domain of dyskerin (21, 24, 25). Whereas the effects for some disease mutants have been evaluated for effects on protein-protein and protein-RNA interactions (26–29), the effects of Dyskeratosis congenita mutations on the pseudouridylation activity of H/ACA snoRNPs have not been addressed.

Here, we report high-resolution cryo-EM structures of endogenous *Trichoplusia ni* H/ACA snoRNPs that provide a structural rationale to understand the effects of Dyskeratosis congenita mutations and reveal molecular details underlying communication between H/ACA snoRNP protomers. We show that interactions between protomers in the H/ACA snoRNP coordinate pseudouridylation activity mediated by each snoRNA hairpin.

Mutations within these interfaces affect the activity in each protomer asymmetrically, uncoupling activity between the two halves of the H/ACA snoRNP. We find that several uncharacterized Dyskeratosis mutations within Cbf5 decrease pseudouridylation activity and assembly of the H/ACA snoRNP. Moreover, we uncover structural coordination between the Cbf5 N-terminal extensions, the Nop10 C-terminus and the RNA interaction with Nhp2 that regulates pseudouridylation activity. Together, our work provides critical insight into the mechanistic coordination of pseudouridine formation across the two H/ACA snoRNP protomers and the molecular defects of Dyskeratosis congenita substitutions in dyskerin.

## Results

### Purification of the insect H/ACA snoRNP

We isolated the H/ACA snoRNP complex from *Trichoplusia ni* high five™ cells (hereafter referred to as H/ACA snoRNP) after heterologous expression and purification of the human HUSH complex (**Fig. S1**). Interestingly, the H/ACA snoRNP complex co-purified with human HUSH after initial affinity pulldown of FLAG-Tasor. However, human HUSH and the H/ACA snoRNP eluted separately from size exclusion chromatography following overnight incubation on ice, suggesting that human HUSH and the H/ACA snoRNP do not directly interact and may instead be linked by some labile component. Because HUSH and the H/ACA snoRNP both bind to RNA (30, 31), we hypothesize that some common RNA substrate transiently links the two complexes during affinity purification, but after extended incubation, this RNA species may degrade or dissociate, causing the HUSH complex and the H/ACA snoRNP to separate.

We used mass spectrometry to verify the identity of the purified H/ACA snoRNP components and positively identified peptides from *T. ni.* Cbf5, Nhp2, Gar1 and Nop10 (**Fig. 1c, Table S1**). We then isolated and sequenced the RNA molecules that co-purified with the H/ACA snoRNP complex (**Table S2, Methods**). Hundreds of *Trichoplusia ni* genes were identified in the copurified RNAs with many of the most prominent RNAs encoding ribosomal proteins (**Table S2**). Because HUSH binds to retrotransposon RNA, we aligned the co-purified RNAs to *Trichoplusia ni* repeats annotated in repeat masker (32). Many of the top repeats identified are lines, sines and helitrons, agreeing with the known binding preference of HUSH for retrotransposon RNA (**Table S3**). Unsurprisingly, we also identified many ribosomal RNAs (rRNA), which are known substrates of the H/ACA snoRNP. Because H/ACA snoRNAs are not well annotated in the *Trichoplusia ni* genome, we assembled contigs for the most common RNA species in the purified H/ACA snoRNA sample **(Table S4)**. Many of the most abundant sequences contain clear features conserved in eukaryotic H/ACA snoRNAs including a two stem-loop architecture and sequence motifs corresponding to the H and ACA boxes (**Fig. 1a, d, Fig. S2).** Therefore, we conclude that our sample contains a mixture of endogenous H/ACA snoRNPs from *Trichoplusia ni* that were copurified with human HUSH, likely through shared RNA substrates.

### Structure of the endogenous H/ACA snoRNP from *Trichoplusia ni*

Because no structure of a fully assembled, catalytically active eukaryotic H/ACA snoRNP has been reported, we used cryo-EM to determine several structures of the endogenous H/ACA snoRNPs that copurified with human HUSH. These structures represent different conformations and assembly states of the endogenous H/ACA snoRNP complex **(Fig. S3, S4, Table S5)**. Class I was resolved to the highest resolution, had the clearest cryo-EM density for all subunits, and was used for all subsequent analysis. The resulting 2.92Å cryo-EM reconstruction shows density for the 5′ and 3′ protomers of the H/ACA snoRNP complex, each containing Nhp2, Cbf5, Nop10, Gar1, and one stem loop of the snoRNA (**Fig. 1e)**. We used Alphafold 2 (33) predictions for each snoRNP subunit and the H/ACA component of telomerase RNA (19) as initial models to build a complete structure of the fully assembled H/ACA snoRNP complex. Our structure closely resembles structures of the telomerase H/ACA snoRNP (18–20), except that the endogenous *Tn.* H/ACA snoRNP lacks the telomerase specific subunit TCAB1 and shows clear density for the upper and lower portions of both snoRNA stem loops (**Fig. S5**).

### Interactions between protomers are important for snoRNP assembly and activity

It is not known whether interactions between protomers in the H/ACA snoRNP are functionally important for snoRNA binding, substrate RNA binding, or pseudouridine formation. Interactions between the 5′ and 3′ protomers of the H/ACA snoRNP are primarily mediated by three sets of protein-protein interactions (**Fig. 2a**). The first inter-protomer contact is mediated by a network of charge-charge interactions between 3′ Cbf5 E176, 5′ Nop10 R51 and 5′ Nhp2 E102 (Cbf5 E144, Nop10 R51 and Nhp2 E95 in yeast) (**Fig. 2b, Table S6, Figure S6**). To assess the functional role of this interaction, we generated a Cbf5 (E144R) substitution but were not able to express this variant in *E. coli*, suggesting that this interface is important for protein stability.

**Figure 2.**
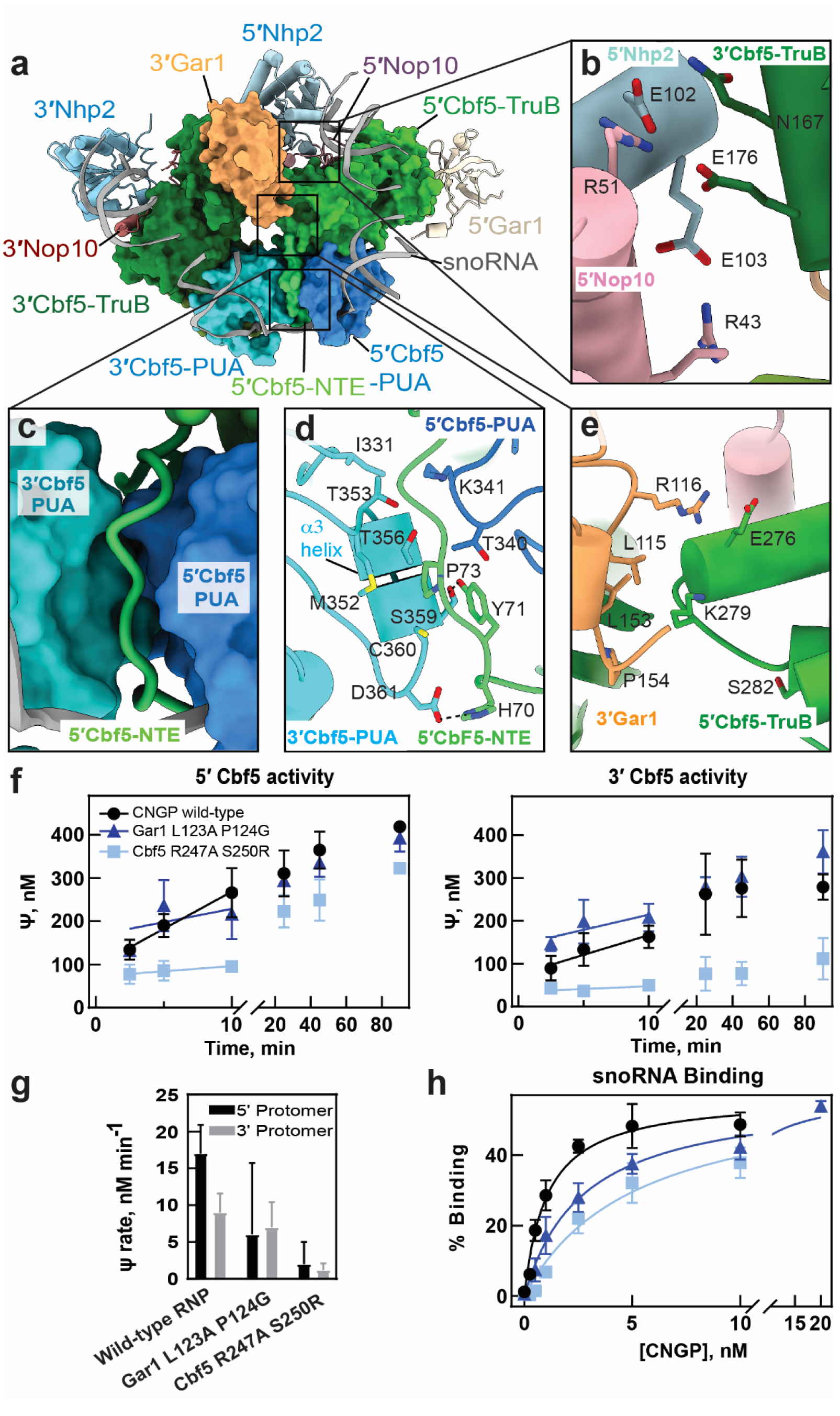
Protein-protein contacts at the dimeric interface coordinate the structure and function of H/ACA snoRNPs. **(a)** Cryo-EM reconstruction of class I snoRNP complex - 5′, 3′ Cbf5 and 3′ Gar1 are highlighted as surface representation in the class I H/ACA snoRNP while other subunits are shown as ribbon representation. The subunits are colored as indicated **(b)** Close-up view of the interface formed at 5’Nop1-5’Nhp2-3’Cbf5. **(c)** Interactions made by 5′-NTE domain with 5′Cbf5-PUA and 3′Cbf5-PUA domains. **(d)** Interaction of 3′-PUA domain with 5′Cbf5 is shown where D361 and S359 are interacting with H67 and Y68 respectively (**e**) Interaction between 5′ and 3′ Gar1 is shown, wherein, K279 of 5′Cbf5-TruB contacts L153 P154 of 3′Gar1. **(f)** Tritium release assays measurement of pseudouridine formation by the snR34 snoRNP modifying a substrate RNA targeted by the 5′ hairpin or 3′ hairpin complex. The yeast Cbf5 variants R247A S250G and Gar1 L123A P124G were functionally characterized relative to wild type. **(g)** Initial velocities of pseudouridylation measured for 5’/3’ hairpin complex are indicated for the yeast Cbf5 variants. **(h)** Filter binding assays determined the binding of the Cbf5-Nop10-Gar1-Nhp2 (CNGP) protein complex to snR34 guide RNA.

The second interaction occurs between a section of the 5′ Cbf5 N-terminal extension (NTE) spanning residues 70-75, and ɑ3 of the 3′ Cbf5 PUA domain (**Fig. 2c, d**). Although this interaction is primarily mediated by the NTE of 5′ Cbf5, the 5′ PUA domain also contacts the 3′ PUA domain using T340 and K341 (**Fig. 2d**). In the middle of the interface, 3′ Cbf5 M352, C360 and T356 comprise a small hydrophobic core with 5′ Cbf5 Y71, P73 and T340 that is ringed by polar interactions. In particular, 5′ Cbf5 H70 and Y71 interact with 3′ Cbf5 S359 and D361. Notably, mutations in human dyskerin at positions corresponding residues H70, M352 and D361 (H68Q, M350T/I, D359N) cause Dyskeratosis congenita (**Table S6, Figure S6, and S7**) (21).

To assess the functional relevance of these interactions, we designed two yeast Cbf5 variants containing either a Cbf5 (H38A, Y39A) double mutant (H70 and Y71 in *T. ni*) or a Cbf5 M320A, S327A, D329A triple mutant (M352, S359 and D361 in *T. ni*). However, the Cbf5 (M320A, S327A, D329A) variant failed to express in *E. coli* despite several attempts, while the Cbf5 (H38A, Y39A) variant expressed but formed insoluble aggregates that precluded purification. These observations suggest that the PUA-NTE interface is critical for Cbf5 stability, and that disruption of this interface makes the protein prone to aggregation during expression, potentially explaining why mutations in this interface can cause Dyskeratosis congenita.

The third interaction occurs between 3′ Gar1 and the catalytic TruB domain of 5′ Cbf5 (**Fig. 2a, e**). In this interface, the aliphatic portion of 5′ Cbf5 K279 (R247 in yeast) contacts 3′ Gar1 L115 and L153 near P15 (
⏢85⏢, L123 and P124 in yeast). In addition, 3′ Gar1 R116 (N86 in yeast) contacts 5′ Cbf5 E276 (D244 in yeast), but Gar1 R116 is not conserved in yeast (**Fig. 2e, Fig. S6**). Notably, this cross-protomer interface is close to Cbf5 S282 (S280 in human and S250 in yeast) which is mutated (21) (22). It is possible that S280R disease mutations could disrupt the local protein structure near the 5′ Cbf5 - 3′ Gar1 interface, potentially causing defects in snoRNP function. To investigate the functional role of the contact between 5′ Cbf5 and the 3′ Gar1, we reconstituted two variants of the yeast H/ACA complex containing Cbf5 (R247A, S250R) or Gar1 (L123A, P124G)⏢We utilized an *in vitro* reconstitution of the *S*. *cerevisiae* H/ACA snoRNP to ⏢(12, 35, 36)⏢. The Gar1 (L123A, P124G) variant affected the activity of each protomer asymmetrically, decreasing the initial velocity of pseudouridylation by about 2-fold for the 5′ hairpin while pseudouridylation targeted by the 3′ hairpin of snR34 was not impaired (**Fig. 2f-g, Table S7**). The Cbf5 (R247A, S250R) variant also reduced pseudouridine formation in both protomers to different extents, corresponding to an 8-fold reduction in the 5′ hairpin and a 4-fold reduction in the 3′ hairpin compared to WT (**Fig. 2f-g, Table S7**). To understand if these defects in pseudouridine formation were due to compromised snoRNA binding, we determined the affinity of Cbf5-Nop10-Gar1-Nhp2 complexes (CNGP) to the snR34 H/ACA snoRNA. The affinity of snR34 snoRNA for Cbf5-Nop10-Gar1-Nhp2 complexes containing Gar1 (L123, P124A) or Cbf5 (R247A, S250R) were two-fold and five-fold lower than for WT complexes (**Fig. 2h, Fig. S8, and Table S8**). However, the pseudouridylation assays were conducted under saturating protein concentrations. Therefore, the small differences in snoRNA affinity for Gar1 (L123, P124A) and Cbf5 (R247A, S250R) probably do not drive the pseudouridylation defects for these mutants. The asymmetric defects caused by mutations in the 5′ Cbf5 – 3 ′Gar1 interface reduced activity in the 5′ protomer more than in the 3′ protomer, suggesting that inter-protomer contacts coordinate the activities of each protomer in the H/ACA snoRNP.

Furthermore, disruption of these inter-protomer contacts decouples the activity of each protomer and compromises pseudouridylation.

### Functional implications of Dyskeratosis congenita mutations

Cbf5/dykerin, Nhp2, and Nop10 harbor a large number Dyskeratosis congenita mutations (21) with most mutations occurring within Cbf5/dyskerin. However, the functional effects of Dyskeratosis congenita mutations on pseudouridine formation have not been directly investigated. Most residues mutated in Dyskeratosis congenita patients are conserved in the *T. ni* H/ACA snoRNP and are concentrated in the PUA domains and the NTE elements as previously described. A few mutations also fall in the catalytic TruB domain of Cbf5, as well as in Nhp2 and Nop10 (**Fig. 3a-d, Fig. S6, Fig. S7 a-c, Table S6**) (21, 34).

**Figure 3.**
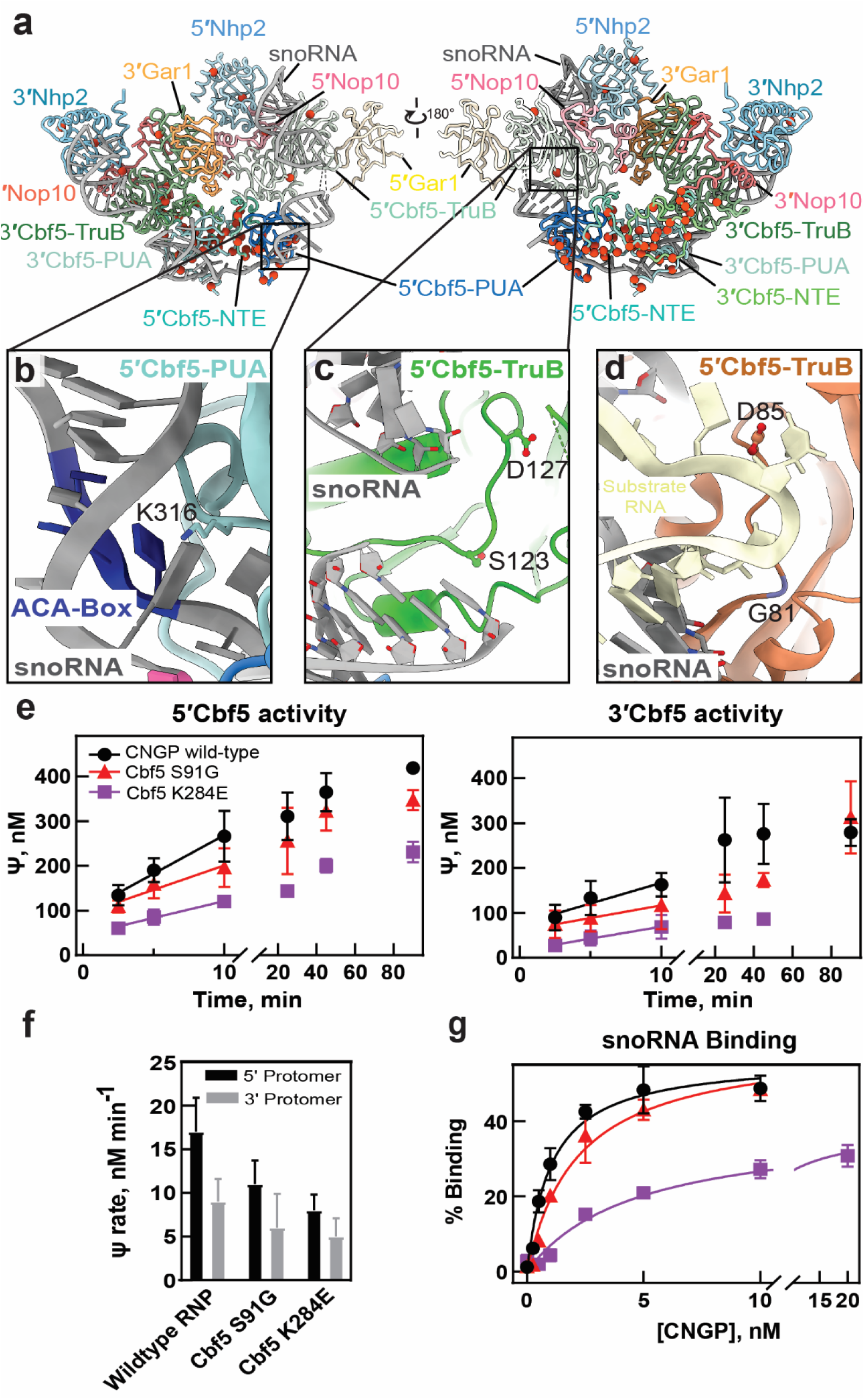
Influence of Dyskeratosis congenita (DC) patient mutations of Cbf5/Dyskerin on snoRNA binding and its pseudouridylation activity. **(a)** DC mutations are mapped on the insect snoRNP complex. Mutations are shown as red spheres. **(b)** Within the Cbf5 PUA domain, DC mutation residue K316 is shown in contact with snoRNA. **(c)** The catalytic center residues S120 and D124 (part of the SGLTD motif) of 5’Cbf5-TruB, implicated in DC are highlighted in the close-up view. **(d)** Comparison of (**c**) to the active site of archaeal H/ACA sRNPs with substrate RNA bound (PDB ID: 3HAY: D85, the catalytic residue of archaeal snoRNP complex. Residue G81 corresponding to insect S120 is shown in close contact with substrate RNA. **(e)** H/ACA snoRNPs reconstituted with yeast Cbf5 variants harboring the substitutions K284E or S91G were analyzed for pseudouridine formation in a substrate RNA recognized by the 5′ hairpin of snR34 as well as in another substrate RNA targeted by the 3′ hairpin of snR34 **(f)** Initial velocities of pseudouridylation measured for 5’/3’ hairpin complex are indicated for the yeast Cbf5 variants - K284E or S91G. **(g)**. Filter binding assays detect the binding of the snR34 H/ACA snoRNA to the Cbf5-Nop1-Gar1-Nhp2 (CNGP) protein complex.

To assess the mechanistic effects of Dyskeratosis congenita substitutions in the PUA domain, we selected K316 (corresponding to K284 in *S. cerevisiae* and K314 in humans) which is mutated in Dyskeratosis congenita patients from multiple families (21). K316 directly interacts with the H/ACA snoRNA in each PUA domain and is located directly adjacent to the ACA box in the 3′ PUA domain (**Fig. 3b**), suggesting that this residue may be important for the proper incorporation of snoRNA into the H/ACA snoRNP. We generated a K284E charge reversal mutation in *Sc*. Cbf5 to disrupt its interaction with the H/ACA RNA. Pseudouridine formation mediated by the 5′ and 3′ hairpins of the snR34 snoRNA were approximately two-fold slower in the K284E mutant compared to WT (**Fig. 3e-f, Table S7**). To understand how the K284E mutation affects snoRNP assembly, we quantified snR34 H/ACA snoRNA binding to the yeast Cbf5(K284E)-Nop10-Gar1-Nhp2 complex. Binding of the Cbf5 (K284E) complex is reduced about 4-fold to the full-length snR34 as well as the individual snR34 5′ hairpin, but 11-fold to the 3′ hairpin (**Fig. 3g, Fig. S8, Table S8**). This indicates the K284E mutation may preferentially disrupt binding to the 3′ hairpin, which contains the ACA box. Importantly, the pseudouridylation assays (**Fig. 3 e-f**) were conducted under saturating protein concentrations, so the defects in pseudouridylation in the Cbf5(K284E) mutant most likely result from a catalytic defect. This suggests that Dyskeratosis congenita mutations within the PUA domains affect both snoRNA binding and pseudouridine catalysis.

Dyskeratosis mutations also occur in the catalytic TruB domain of Cbf5. Cbf5 S121 is part of the conserved SGTLD motif (21) and is commonly mutated to glycine in patients with Hoyeraal-Hreidarsson syndrome, a severe form of Dyskeratosis congenita. In our structure, Cbf5 S123 (equivalent to S121 in humans) is near the D127 catalytic residue and docks into a hydrophobic pocket in the folded TruB domain (**Fig. 3c**). However, the importance of this residue in pseudouridine formation is not clear since the corresponding residue in archaea (G81) does not interact with the substrate RNA (**Fig. 3d**). Pseudouridine formation mediated by both H/ACA snoRNP hairpins in the yeast Cbf5 (S91G) snoRNP was 1.4-fold lower than in complexes containing WT Cbf5 (**Fig. 3e-f, Table S7**). In addition, the affinity of the Cbf5 (S91G) complex for snR34 snoRNA was reduced by approximately 2-fold compared to WT (**Fig. 3g, Fig. S8, Table S8**). Taken together, these data show that Dyskeratosis congenita associated mutations in the PUA and TruB domains of Cbf5/dyskerin cause H/ACA RNA binding defects and catalytic defects in the H/ACA snoRNP.

### Folding of the Cbf5 NTE is asymmetrically correlated with Nhp2 and Cbf5 activity

During cryo-EM processing we identified four distinct classes of the snoRNP in different conformations and assembly states which revealed coordinated structural transitions that influence coupling between protomers and dyskerin activity. In the fully assembled complex (class I), a pyrimidine at the tip of each stem loop (modeled as C31 and C102) docks into a pocket on Nhp2. In addition, the N-terminal extensions (NTE) of 5′ Cbf5 and 3′ Cbf5 form an intertwined structure that docks onto the back of 3′ Cbf5 and contacts the C-terminal part of 3′ Nop10 (**Fig. 4 a-d**). This arrangement is largely retained in class IV, which lacks 5′ Gar1 (**Fig. S9 a**). However, the absence of 5′ Gar1 coincides with specific unfolding of the 5′ Cbf5 thumb loop, which is mostly structured in 5′ Cbf5 from Class I (**Fig. S9b).** Folding of the 3′ and 5′ NTEs is also consistent with a previous structure of the human telomerase H/ACA snoRNP (18). Interestingly, many residues within the Cbf5 NTEs are mutated in Dyskeratosis congenita (34) but it is not known how mutations in the NTE may affect the pseudouridylation activity of Cbf5.

**Figure 4.**
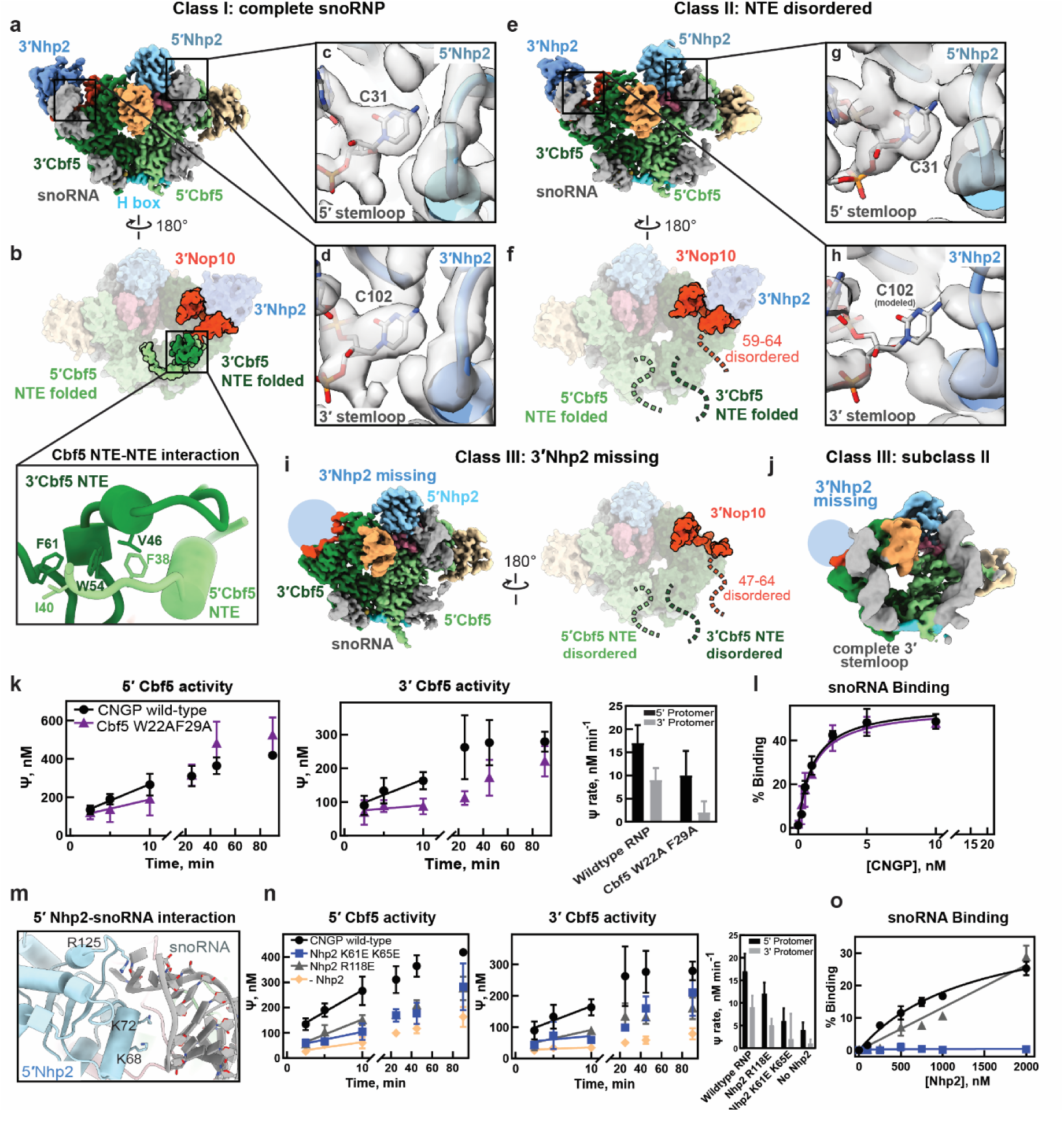
**(a-b)** Cryo-EM structure of class I H/ACA snoRNP, front (a) and back views are shown here (b) with well folded NTE of 5′ and 3′ Cbf5 and N-terminal part of 3′ Nop10. Close-up view of the interaction of hydrophobic interactions between the NTE elements of 5′ and 3′ Cbf5 (inset). **(c-d)** Close-up views show the interaction of 5′ and 3′ stem-loops with the 5′ and 3′ Nhp2 (c and d) respectively. **(e-f)** Cryo-EM structure of class II H/ACA snoRNP, Front (e) and back views are shown here (f) with disordered NTE of 5′ and 3′ Cbf5 and N-terminal part of 3′ Nop10 (residues 59-64). **(g-h)** Close-up views show the interaction of 5′ and 3′ stem-loops with the 5′ and 3′ Nhp2 (g and h) respectively. **(i)** Cryo-EM structure of the 3′ Nhp2 missing class III snoRNP (both front and back views are represented here). The NTE of 5′ and 3′ Cbf5 and N-terminus (47-64 residues) of 5’ Nop10 which are disordered in class III are indicated. **(j)** subclass II of class III (Nhp2 missing class) showing complete 3′ stem-loop. **(k)** Tritium release assay measurement of pseudouridine formation for substrate RNA targeted by 5’ or 3’ hairpin of snR34 including the yeast Cbf5 W22A F29A substitutions. Also shown are the initial velocities measured in the assay. **(l)** Filter binding assays detect the binding of the snR34 H/ACA snoRNA to the Cbf5-Nop1-Gar1-Nhp2 (CNGP) protein complex containing the yeast Cbf5 W22A F29A substitutions. **(m)** Charged residues of 5′ Nhp2 (K68, K72 and R125) interaction with the stem-loop of 5′ H/ACA snoRNA. **(n)** Tritium release assay measurement of pseudouridine formation for substrate RNA targeted by 5’ or 3’ hairpin of snR34 including the yeast Nhp2 R118E and K61E K65E substitutions and in the absence of Nhp2. Also shown are the initial velocities measured in the assay. **(o)** Filter assay binding to quantify the interaction between Nhp2 wt and variants with snR34 H/ACA snoRNA.

Further classification of our cryo-EM dataset revealed a second class of fully assembled H/ACA snoRNP molecules (class II) that differs from class I in the structure of the Cbf5 NTEs. In class II, the NTE of both Cbf5 monomers and C-terminal end of 3′ Nop10 (59–64) are completely disordered, and density for C102 in the pocket of 3′ Nhp2 is missing (**Fig. 4 e-h**). We observed similar structural changes in Class III where 3′ Nhp2 is absent. In this structure, density for the upper portion of the 3′ stem loop is completely missing whereas the 5′ stem loop binds 5′ Nhp2 in a similar conformation seen in the other classes (**Fig. 4i**). In addition, the Cbf5 NTEs and a longer C-terminal portion of Nop10 (47–64) are completely unfolded in a similar manner as in Class II (**Fig. 4i**).

Because Nhp2 makes critical contacts with the snoRNA, we performed deep classification of Class III to understand how the 3′ RNA stem loop is affected by the loss of Nhp2. Many subclasses of Class III had no interpretable density for the upper part of the 3′ stem loop (**Fig. S9c**) indicating that without Nhp2, the upper portion of the 3′ stem loop is unfolded or highly mobile. We identified one class (subclass II) that contained a complete 3′ stem loop in the absence of Nhp2 **(Fig. 4j)**. However, the 3′ stem loop is moved away from the rest of the snoRNP complex and is not positioned to direct a substrate RNA into the Cbf5 active site (**Fig. S9d).** This arrangement is similar to a structure of the archaeal H/ACA sRNP lacking the Nhp2 homolog L7Ae (17). Together, these observations show that folding of the Cbf5 NTEs and the 3′ Nop10 C-terminus is correlated with the 3′ stem loop binding to 3′ Nhp2.

To understand the functional importance of the coordinated structural transitions observed for the Cbf5 NTEs, we disrupted interactions within the NTEs and determined effects on pseudouridylation in the 3′ and 5′ protomer. The 5′ and 3′ Cbf5 NTEs contact each other using an extended set of hydrophobic interactions including residues F38, I40 of 5’ Cbf5 and V46, W54, F61 of the 3′ Cbf5 (**Fig. 4b, inset**). To understand how disruption of NTE folding impacts Cbf5 activity, we reconstituted *S. cerevisiae* H/ACA snoRNP complexes containing either WT Cbf5 or a Cbf5 variant containing a W22A and F29A double mutation (corresponding to W54 and F61 in *T. ni*) (12, 35, 36). The Cbf5 (W22A, F29A) mutant reduced pseudouridine formation in the 5′ protomer by 2-fold, but reduced activity in the 3′ protomer by 4-fold compared to WT complexes (**Fig. 4k, Table S7**).However, the affinity of the Cbf5-Nop10-Gar1-Nhp2 complex for the snR34 H/ACA snoRNA and the individual hairpins of snR34 were not affected by the Cbf5 (W22A, F29A) mutations (**Fig. 4l, Fig. S8, Table S8**). Therefore, disruption of the Cbf5 NTEs primarily affects the catalytic activity of Cbf5. Interestingly, the effect of NTE disruption on Cbf5 activity is not symmetrical as pseudouridylation guided by the 3′ hairpin was more affected than pseudouridylation directed by the 5′ hairpin. Because the folded NTEs dock onto the 3′ protomer, it seems likely that disruption of NTE folding would primarily affect the activity of 3′ Cbf5. This observation highlights the importance of this inter-protomer interaction for H/ACA snoRNP function and indicates that the activity of the two H/ACA snoRNP protomers is coordinated.

Next, we asked how the interaction between Nhp2 and the H/ACA snoRNA affects H/ACA snoRNP function. In Class II and III the interaction between Nhp2 and the 3′ snoRNA hairpin is correlated with folding of the Cbf5 NTEs, which asymmetrically affects activity in the 3′ protomer (**Fig. 4k**). We identified two additional electrostatic interactions between Nhp2 and the H/ACA snoRNA. Nhp2 residues K68 and K72, directly interact with the RNA backbone and R125 contacts the tip of each hairpin (**Fig. 4m**). We substituted these residues in yeast Nhp2 (K61, K65, and R118, respectively) with glutamate to disrupt the interaction between Nhp2 and the H/ACA snoRNA **(Table S9, S10)**. The Nhp2 (K61E, K65E) variant strongly reduced pseudouridylation mediated by both protomers by 3- to 4-fold. This is similar to the defect observed for snoRNP reconstituted without any Nhp2 (**Fig. 4n, Table S7**). In addition, the Nhp2 R118E variant reduced pseudouridylation in both protomers by 1.5-fold (**Fig. 4n, Table S7**). To confirm that these substitutions disrupt interactions with the snoRNA we measured affinity of the different Nhp2 variants to the snR34 H/ACA snoRNA. Nhp2 (K61E, K65E) completely abolished its interaction with RNA, and Nhp2 (R118E) strongly reduced binding such that a reliable dissociation constant could not be measured (**Fig. 4o**). These observations suggest that the Nhp2-RNA interaction is critical for pseudouridylation and that Nhp2 mutations affect Cbf5 activity equally in both protomers.

## Discussion

H/ACA snoRNPs are biologically and medically significant as they facilitate critical cellular processes such as ribosome biogenesis and mammalian telomere maintenance. However, a detailed understanding of coordination of pseudouridylation activity across the two protomers of eukaryotic, dimeric H/ACA snoRNPs remains lacking (13, 14, 17, 37, 38). The insect H/ACA snoRNP structure reported here provides for the first time insight into the pseudouridylation-active, eukaryotic complex and allows for the systematic analysis of the structure-function relationship of the protomer interactions for RNA pseudouridylation including the impact of Dyskeratosis congenita disease mutations. Interestingly, we identify critical interaction sites between the two protomers that are important for coordinating the pseudouridylation activity in both protomers. In addition, we describe for the first time that Dyskeratosis congenita mutations can directly impact pseudouridine formation.

### Coordination of H/ACA snoRNP activity across the two protomers

In addition to the known interaction between the PUA domains and NTEs of the 5′ and 3′ Cbf5, we also observe protein-protein interactions between the 3′ Gar1 and the 5′ Cbf5 proteins as well as between the 3′ Cbf5, 5′ Nop10 and 5′ Nhp2 proteins (Fig. 2). Notably, the disruption of the contact between the 3′ Gar1 and the 5′ Cbf5 asymmetrically affects pseudouridylation reducing the activity of the 5′ protomer more than of the 3′ protomer. Presumably, the 3′ Gar1 - 5′ Cbf5 interaction affects the active site of 5′ Cbf5, which is in close vicinity. Specifically, mutation of Cbf5 residues R247 and S250 may disrupt the local structure of 5′ Cbf5 and potentially propagate that disruption to helix 7 whose N-terminus lies behind the active site. Surprisingly, the mutations in Gar1 and Cbf5 also impair binding of the H/ACA snoRNA to the proteins, possibly due to an overall mispositioning of the two promoters which may indirectly affect the binding of H/ACA snoRNA. In conclusion, the differential effects of disrupting the 3′ Gar1 - 5′ Cbf5 interaction on the modification activity by both protomers suggest an unexpected role for this protein-protein interaction in enhancing the activity of the 5′ protomer. Additionally, we identify a contact between the two protomers mediated by Cbf5 NTEs. Upon disrupting the folding of the NTEs and their interaction in yeast Cbf5 W22A F29A, pseudouridylation is significantly more affected in the 3′ protomer than in the 5′ protomer. The position of this interaction is relatively closer to the active site of the 3′ Cbf5 than the 5′ Cbf5 as the folded NTEs are docked onto the 3′ protomer. This suggests that allosteric effects of unfolding the NTEs differentially influence the two catalytic centers.

Together, these findings demonstrate that the 3′ Gar1 - 5′ Cbf5 interaction enhances activity in the 5′ protomer, while the Cbf5 NTE interaction augments the pseudouridylation activity in the 3′ protomer. Therefore, the protomer-protomer interactions are increasing the RNA modification activity mediated by both H/ACA snoRNA hairpins. This finding explains for the first time mechanistically why eukaryotic H/ACA snoRNPs benefit from containing two hairpins and two sets of proteins.

Such an asymmetric assembly of enzymes is highly unusual as most multimeric enzymes display symmetry where inter-protomer interactions allosterically regulate both active sites (39). The asymmetry observed for H/ACA snoRNPs likely stems from the interaction of the subunits with the single, two-hairpin H/ACA snoRNA giving rise to unique asymmetric interactions influencing the two active sites in Cbf5/dyskerin. Interestingly, the homodimeric human ADAR2 enzyme responsible for RNA deamination is also characterized by an asymmetric assembly with functional implications. Here, only one catalytic site is active whereas interaction with the second catalytic domain increases the RNA deamination velocity of the first catalytic domain (40). With the rapid advancement in structure determination and prediction, additional asymmetric enzymes may be discovered, and it will be interesting to study the mechanistic and evolutionary features of such enzyme dimerization.

### Mechanistic insights into Dyskeratosis congenita mutations

Mutations in all domains of Cbf5/dyskerin including in the catalytic domain as well as Nop10 and Nhp2 (and other proteins) cause the inherited bone-marrow failure syndrome Dyskeratosis congenita. While the molecular cause of Dyskeratosis congenita is complex, the effects of several mutations have been analyzed with respect to protein-RNA and protein-protein interactions (26, 28, 29, 41–43). However, it remains unclear whether Dyskeratosis congenita mutations can also directly impair the pseudouridylation activity of H/ACA snoRNPs and the interaction with H/ACA snoRNAs guiding these modifications. Here, we demonstrate that dyskerin mutations in the catalytic domain (S121G and S280R in humans, S91 and S250 in yeast) as well as in the PUA domain (K314R in humans, K284 in yeast) cause a strong decrease in pseudouridine formation mediated by both protomers. Telomerase stability and function does not depend on the catalytic activity of dyskerin, and the same is true for the 18S rRNA folding activity of the essential U17/snR30 H/ACA snoRNA. Therefore, the mutations in Cbf5 characterized here most likely contribute to the disease’s symptoms by decreasing modification of ribosomal RNA in Dyskeratosis congenita patients which in turn results in impaired ribosome biogenesis and ribosome function (22, 44, 45). As such, our findings provide additional evidence that Dyskeratosis congenita is caused by both telomerase and ribosome biogenesis defects.

### Functional interplay between Cbf5’s NTEs, Nop10 and Nhp2 in the 3′ protomer

For the first time, we describe coordinated conformations of the two Cbf5 NTEs and the 3′ Nop10 C-terminus on the backside of the 3′ Cbf5 that are further correlated with the interaction of the 3′ Nhp2 with H/ACA snoRNA. Specifically, in the Class I conformational state, the Cbf5 NTEs interact and contact the C-terminus of Nop10 while Nhp2 contacts the H/ACA snoRNA. However in Class II and IV, all these interactions are lost simultaneously, and mutations disrupting these interactions between the Cbf5 NTEs or between Nhp2 and snoRNA result in decreased pseudouridylation activity of the 3′ protomer. Accordingly, the conformational state lacking these interactions can be described as less active. This observation suggests a potential mechanism for regulating the activity of H/ACA snoRNPs by disrupting the Cbf5 NTE interactions.

These observations also further underline the functional importance of Nhp2’s interaction with the H/ACA snoRNA regardless of Nhp2’s lower affinity and specificity compared to the archaeal L7Ae homolog (46). Despite an intact interaction between Nhp2 and Nop10, the pseudouridylation activity is strongly reduced for two Nhp2 variants impaired in RNA binding. Thus, the interaction of Nhp2 with the tip of the H/ACA snoRNA hairpin is critical for correctly positioning the H/ACA snoRNA relative to the catalytic site of Cbf5 (12), but also for stabilizing the interaction of Nop10’s C-terminus with the NTEs of Cbf5. Together, our structural and functional characterization of catalytically active H/ACA snoRNPs thus reveal for the first time an intricate conformational coordination across the entire ribonucleoprotein complex.

### Conclusions and outstanding questions

Determining the structure of a eukaryotic, catalytically active H/ACA snoRNP allowed us to dissect the structure-function relationship of critical protein-protein and protein-RNA interactions within and across the two protomers characteristic for eukaryotic (but not archaeal) H/ACA snoRNPs. Thereby, we uncovered critical points of coordination between the two protomers that enhance either the activity of the 3′ protomer (in case of the 3′ Gar1 - 5′ Cbf5 interaction) or the activity of the 5′ protomer (in the case of the Cbf5 NTEs interaction). Evidently, eukaryotic H/ACA snoRNPs benefit from being assembled on two hairpins in H/ACA snoRNA rather than a single hairpin as found in some archaeal H/ACA sRNAs. On the other hand, our structure-function analysis raises the question why eukaryotic H/ACA snoRNAs are not comprised of more than two hairpins. From a structural perspective, there is space to add a third protomer to the 3′ end of the H/ACA snoRNA which could possibly further enhance the pseudouridylation activity in particular of the middle protomer. We speculate that the demands of other interactions in the cell, in particular the dense nucleolus, limit the size and complexity of eukaryotic H/ACA snoRNPs. As the structure of insect H/ACA snoRNP contains several different H/ACA snoRNAs, we were unable to determine a homogeneous structure of insect H/ACA snoRNP bound to substrate RNA. Therefore, it remains to be investigated how large RNAs such as ribosomal RNA with multiple pseudouridylation sites interact with the two pseudouridylation pockets in eukaryotic H/ACA snoRNP. In conclusion, our study provides detailed insight into the sophisticated architecture and function of H/ACA snoRNPs that explains their role in ribosome biogenesis, telomerase maintenance and other functions in healthy cells as well as the molecular defects underlying Dyskeratosis congenita.

## Materials and Methods

A complete description of H/ACA snoRNP purification, structure determination, pseudouridylation assays, H/ACA snoRNA binding assays and other methods can be found in the Supporting Information.

Cryo-EM maps and structural models have been deposited into the Protein Data Bank. Class I (XXXX), Class II (XXXX), Class III (XXXX), Class IV (XXXX).

## Acknowledgements and Funding Sources

Research in the Kothe lab was supported by the Canadian Institutes for Health Research (CIHR, Project Grant 437623). Research in the Worden lab was supported by National Institute of General Medical Sciences (NIGMS, 5R35GM147261).

