## Supplemental Information for "Interprotomer Communication and Functional Asymmetry in H/ACA snoRNPs"

1.1 Materials and Methods
1.2 Supplemental Figures
1.3 Tables

### 1.1 Material and Methods

#### Materials

[C5-$^3$H]-UTP was purchased from Moravek Biochemicals. [γ-P32]-ATP was purchased from Revvity. All chromatography materials are from GE Healthcare. Oligonucleotides were ordered from Integrated DNA Technologies (IDT), and synthesized plasmids were ordered from Genewiz. All other chemicals and enzymes were purchased from Fisher Scientific.

#### Cloning, baculovirus generation and Insect cell protein expression of Hush complex

The hush subunits (flag TASOR, MPP8, Morc2, and Periphilin) were cloned into a co-expression vector for insect cell expression using the biGBac vector system described previously (1). Briefly, the four HUSH subunits (flag-TASOR, MPP8, Periphilin and Morc2) were cloned individually into pLib. Then each gene expression cassette containing each protein was amplified with the designated gibson assembly primers from the BigBac system and individually cloned into pBIG1a-d (pBIG1a: flag-TASOR, pBIG1b: MPP8, pBIG1C: periphilin, pBIG1d: MORC2). In the second step, each pBIG1 construct was digested with Pme1 and the inserts containing each gene expression cassette were cloned into to the pBig2 plasmid creating the pBIG2:flag-TASOR, MPP8, MORC2, Periphilin co-expression construct. The clones were confirmed using restriction digestion (PacI), junction PCR and DNA sequencing.

For bacmid generation, DH10Bac™ cells were transformed with the Hush co-expression construct and plated on the bacmid selection LB agar plates. Bacmid was isolated the P0 baculovirus stock was prepped by transfecting of 2.5ml of SF9 insect cells with 5ul of bacmid using Promega FuGENE HD transfection reagent. Sf9 insect cells were incubated in a 24-well cell culture block on a shaker (27°C, 300 rpm, 5 days). The cells were spun down at

2000 rpm for 15 min and supernatant was collected as P0. Subsequently, P1 (40ml) and P2 (100ml) baculovirus were generated by incubation of ~2.5 million cells (SF9)/ml culture with 1% and 2.5% P0 and P1 respectively on a shaker (27°C, 200 rpm, 2 days). The protein was expressed by addition of 1.2% P2 virus stock to 1L of high five cells and incubated for 4 days on a shaker (27°C, 200 rpm). The cells were pelleted (3000 rpm, 15 min.) and flash frozen in liquid N2 and stored at -80 °C.

**Human hush and insect snoRNP complex co-purification**

The frozen pellet was thawed, resuspended in the lysis Buffer (30mM HEPES pH 7.9, 300mM NaCl, 10%Glycerol) and sonicated with stirring continuously in ice water bath for 3 cycles of 40% Amplitude, 5 seconds ON and 55 seconds OFF for 4 mininutes total time. After sonication, the cell extract was clarified by centrifugation and the clarified supernatant was incubated with anti-DYKDDDK resin (genscript) pre-equilibrated with lysis buffer for 1hr on rocker at 4 °C in a 50ml conical tube. After 1 hr, the affinity resin was spun down at 400 rpm for 15 minutes, and the beads were washed 2x with lysis buffer. After washing, the beads were resuspended in 50ml lysis buffer and incubated for another 15 minutes at 4°C. The affinity resin was transfered to a gravity column and washed with another 50ml of lysis buffer. The bound protein was eluted in 1ml fractions using Lysis Buffer supplemented with 0.15 mg/ml anti-flag peptide (Medchem Express - Cat. No.: HY-P0223). The elution fractions were analyzed on 12% SDS-PAGE and the protein containing fractions were pooled together and incubated on ice overnight with fresh addition of 2mM DTT. The next day, the pooled fractions were concentrated and loaded on a Superose 6 gel filtration column pre-equilibrated with lysis buffer + 2mM DTT. The protein was eluted as 2 distinct peaks with the first peak corresponding to hush subunits and the second peak corresponding to the endogenous snoRNP. Pooled fractions were concentrated, flash frozen in LN2 and stored at -80°C.

**Cloning, protein expression and purification of Yeast H/ACA snoRNP complex**

The genes for CBF5, NOP10, GAR1, and NHP2 were cloned as previous (2). CBF5 and GAR1 variants harboring point mutations were generated by Genewiz. Nhp2 variants were created through QuikChange site-directed mutagenesis (Table S6).

All proteins were expressed and purified as previously described (2). Nhp2 proteins are expressed with an N-terminal Histidine tag. Cbf5 proteins contain a C-terminal Histidine tag and are co-expressed alongside wild-type Nop10 from the same plasmid in *E. coli* Lemo21 (DE3) cells (NEB) using 50 μM L-Rhamnose and 1 mM isopropyl β-D-1-thiogalactopyranoside (IPTG). Gar1 wild-type and the variant Gar1 L123A P124G are expressed as GST fusion proteins in *E. coli* Rosseta2 cells (NEB). Nhp2 proteins are purified separately by Nickel-Sepharose chromatography followed by cation exchange chromatography. Cbf5-Nop10 expressing cells are mixed with Gar1-expressing cells to allow for the purification of the Cbf5-Nop10-Gar1 complex by tandem glutathione- and Nickel-Sepharose affinity chromatography.

**Nano Liquid chromatography- Mass spectrometry Analysis**

The snoRNP sample was run on SDS-PAGE and the gel bands were subjected to tryptic digestion using the Tryptic Digestion Kit (Thermo Fisher, PN 89871). Briefly, the gel bands were destained with 200 μL of destaining solution. Reduction was performed by incubating the band in 30 μL of reducing agent solution containing TCEP at 60 °C for 10 minutes with shaking at 700 rpm. Alkylation followed, using 30 μL of alkylating agent solution containing TCEP, incubated at room temperature in the dark for 60 minutes with shaking at 700 rpm.

Proteins were then digested by adding 25.0 µL of digestion solution to each sample, followed by overnight incubation at 30 °C with shaking at 700 rpm on a Thermomixer. The supernatants were dried in a SpeedVac and resuspended in 10 µL of 0.1% TFA in $H_2O$.

Each sample (3 µL) was analyzed on an Orbitrap Eclipse (Thermo Scientific) attached to a Vanquish Neo nano UPLC (Thermo Scientific). Peptides were separated with a nano HPLC column (75 µm x 20 cm, 1.7µm C18, CoAnn Technologies). Peptides were eluted with a linear gradient from 3 % to 25 % of buffer B in 22 min. This was followed by increasing buffer B to 80 % in 4 min and washing with 95 % B in 4 min. The FAIMS Pro source (Thermo Fisher Scientific) was located between the nanoESI source and the mass spectrometer. The selected compensation voltage (CV) was applied (−40V, −55V, and -75V) throughout the LC-MS/MS runs. The column temperature was kept constant at 50 °C using a customized column heater (Phoenix S&T, Chadds Ford, PA). One full MS scan was collected with a scan range of 350 to 1200 m/z, resolution of 120 K, standard maximum injection time, and Auto Gain Control (AGC). Then, a series of MS2 scans were acquired for the most abundant ions from the MS1 scan using an ion trap analyzer. Ions were filtered with charge 2–5. An isolation window of 1.6m/z was used with quadruple isolation mode. Ions were fragmented using higher-energy collisional dissociation (HCD) with a collision energy of 30%.

Proteome Discoverer 3.0 (Thermo Fisher Scientific) was used to process the raw spectra. Database searches were conducted against the human protein database and the SP9 database. Search parameters included carbamidomethylation (+57 Da) of cysteine as a fixed modification and oxidation (+16 Da) of methionine as a variable modification. A maximum of two missed cleavages was allowed. The precursor mass tolerance was set to 10 ppm, and the fragment mass tolerance was set to 0.6 Da.

**snoRNA isolation and sequencing**

We isolated the RNA that co-precipitated with the snoRNP by performing TRIzol (Thermo Fisher Scientific) - isopropanol precipitation following manufacturer's recommendations. We used the SMARTer® smRNA-Seq kit for Illumina (Takara, Cat. 635029) to construct a RNA sequencing library from the extracted RNA. We sequenced the RNA library on the NovaSeq 6000 platform (Illumina) to generate 15 million 100bp paired-end reads. The sequence reads were trimmed for adapter sequences using TrimGalore (v0.6.0, https://zenodo.org/records/7598955). Since small RNA species are often smaller than 200bp (3) we merged the overlapping paired-end reads to create longer single-end reads for alignment with NGmerge (v0.3) (4). Since SMARTer® smRNA-Seq kit artificially polyadenylated RNA molecules during library construction, we removed polyA sequences at 3' end of the reads with fastp tool (v0.23.4) (5). These single-end reads were aligned to the *Trichoplusia ni* (Hi5) (6) reference genome using HISAT2 (v2.2.1) (7). Gene expressions were calculated using StringTie (v2.2.1) (8). We also quantified the number of reads that mapped to annotated small repeats or transposable elements in the Hi5 genome (6) with BEDTools "coverage" function (v2.30.0) (9). Lastly, we utilized the Trinity tool (10) to perform *de novo* reconstruction of the transcriptome to identify unannotated RNA species that were bound to the snoRNP complex.

**Cryo-EM sample preparation and data acquisition**

Briefly, 125nM of the snoRNA complex in 2.4ml of reaction buffer (30mM HEPES pH 7.9, 300mM NaCl, 10% Glycerol, 2mM DTT containing 0.015% glutaraldehyde) was incubated for ice for 20 min and quenched by addition of 1M Tris-HCl to a final concentration of 40mM Tris-HCl. The reaction was concentrated to 200 ul and dialyzed against the buffer (30mM

HEPES pH 7.9, 300mM NaCl, 1mM DTT) and kept overnight at 4 °C. For cryo-EM grids preparation, 4mM of concentrated protein was applied on the glow-discharged (for 30s) Quantafoil Au 1.2/1.3 grids and blotted [2.5s (total time), 0 (Blot force), 1 (Wait time), 1 (Blot time), 6°C and 100% humidity] and flash frozen in liquid ethane.

Cryo-EM data were collected on 300kV Titan Krios electron microscope equipped with a K3 direct electron (Gatan) in super-resolution mode. With a nominal magnification of x105,000 and a pixel size of 0.414 Å. 50 frames were collected for each movie with a total dose of 60 e/A. 15130 movies were collected and processed with CryoSPARC (11).

**Cryo-EM data processing**

The super-resolution movies (15130 movies) were imported to cryoSPARC and motion corrected using the patch motion correction job with the fourier cropping factor set to ½. CTF parameters were computed using the patch CTF estimation job. Blob picker was used to select particles from 1000 micrographs followed by 2D classification to generate templates for template particle picking. Two rounds of 2D classification, followed by an initial ab-initio classification was used to generate the initial 3D classes. After this, 2 rounds of heterogeneous refinement were performed and the best resolved classes particles were then subjected to NU-refinement. Two classes with good map features were subjected to an additional round of heterogeneous refinement that resulted in the final structures of the snoRNP. Heterogeneous refinement of Class I resulted in final classes I and IV. Class I (760,070 particles, 2.92 Å) shows good features for all the protein and RNA components of the snoRNP complex. Class IV (74734 particles, 3.23 Å) is identical to Class I except that a Gar1 subunit is missing. Class I and IV were subjected to deep-EMhancer (12) to sharpen the map.

Heterogeneous refinement of class II resulted in final classes I and III. Final class II (733,250 particles, 2.86 Å) had similar features to final class I except the CTE of each Cbf5 subunit was disordered. To improve the map features of final class II we conducted two rounds of particle-subtraction and local refinement for each SnoRNP protomer. We then created a composite map for final class II by combining the focused maps using the Vop add command in Chimera (14). Final class III (134826 particles, 3.14 Å) is missing density for Nhp2 and the 3′ the stem loop. Class II and III were subjected to deep-EMhancer (12) to sharpen each map.

**Model Building and Refinement**

For each structure, initial models for each snoRNP subunit were generated using Alphafold2 using the sequences of the insect snoRNP subunits - Cbf5, Nop10, Nhp2 and Gar1. RNA from the telomerase H/ACA module (PDB: 8OUF) was used as the initial model for the snoRNA (13). The initial model was docked onto the cryo-EM maps using USCF Chimera (14) and refined using rigid body refinement feature in Phenix (15). This was followed by manual rebuilding of the initial models using Coot (16). This was followed by multiple rounds to iterative model improvement in Phenix (using minimization and local grid search parameters) and Coot.

**In vitro transcription of RNA** - The template for snR34 guide RNA was cloned, transcribed and purified as reported previously (2). Substrate RNAs for the 5’ and 3’ hairpin of snR34, H89 and H90-92, respectively, were generated by *in vitro* transcription in the presence of [C5–$^{3}$H]-UTP using annealed oligonucleotides as a template **(Table S7)**. Excess proteins are removed from the RNA via phenol-chloroform extraction followed by ethanol precipitation of the RNA. Subsequently, free nucleotides are removed using a G-25 sephadex spin column.

To create radiolabeled snR34, the ends were dephosphorylated with Calf intestinal alkaline phosphatase (NEB) and the RNA was purified by phenol-chloroform extraction. [γ-$^{32}$P]-ATP was incubated with T4 Polynucleotide Kinase (Fisher) to label the RNA. Excess nucleotide is removed with a G-25 sephadex spin column.

**Reconstitution of an active yeast H/ACA snoRNP complex** - The H/ACA snoRNP complex is reconstituted from purified snR34, Nhp2, and Cbf5-Nop10-Gar complex as described. In brief, snR34 is first folded by slow cooling from 65°C to room temperature in Reaction Buffer (20 mM HEPESKOH (pH 7.4), 150 mM NaCl, 0.1 mM EDTA, 1.5 mM MgCl2, 10% (v/v) glycerol, 0.75 mM dithiothreitol (DTT)). Subsequently, the proteins are added followed by incubation for 10 min at 30°C such that the H/ACA snoRNP complex is formed. The final concentrations used in pseudouridylation assays are 50 nM for snR34 and 100 nM for each protein to account for the dimeric nature of the complex.

**Nitrocellulose filtration assay -** Nitrocellulose filter binding assays were carried out as previous (2). Binding Reactions were set up in Reaction Buffer with a constant 0.1 nM concentration of [$^{32}$P]-labeled snR34 guide RNA and 0-10 nM H/ACA proteins (Cbf5-Nop10-Gar1 and Nhp2). The reaction was incubated for 10 minutes at 30°C. Afterwards, the reaction was filtered through a 0.2 µm pore nitrocellulose membrane which was washed with 1 ml of cold Reaction Buffer. The membrane was dissolved in 10 ml of scintillation cocktail and then counted using a 4910 TR (Perkin Elmer) liquid scintillation counter. For reactions with Nhp2 wt and variants only, the assay was carried out in an identical manner while omitting the Cbf5-Nop10-Gar1 complex and using an Nhp2 concentration up to 2000 nM. Dissociation constants ($K_D$) were determined by fitting the binding curves to the following hyperbolic function using GraphPad Prism.

$Y = B_{max} \times [P] / (K_D + [P])$ (Equation 1)

with [P] being the protein concentration and $B_{max}$ the maximum binding.

**Tritium release assay -** Tritium release assays were carried out as previous (2). In short, the H/ACA snoRNP complex (50 nM) was reconstituted and radiolabelled substrate RNA was added to the reaction to a final concentration of 500 nM. At different time points, samples were removed and quenched in Norit A charcoal in 0.1 M HCl. Charcoal was removed by centrifugation and filtering through glass wool. The supernatant was counted by liquid scintillation counting (Tricarb 4910 TR, Perkin Elmer). The amount of pseudouridine formation was calculated using the specific activity, and the background (determined in the absence of H/ACA snoRNP) was subtracted. Initial velocities were determined by plotting only the early reaction time points and linear regression where the slope corresponds to the initial velocity.

### 1.2 Supplemental Figures

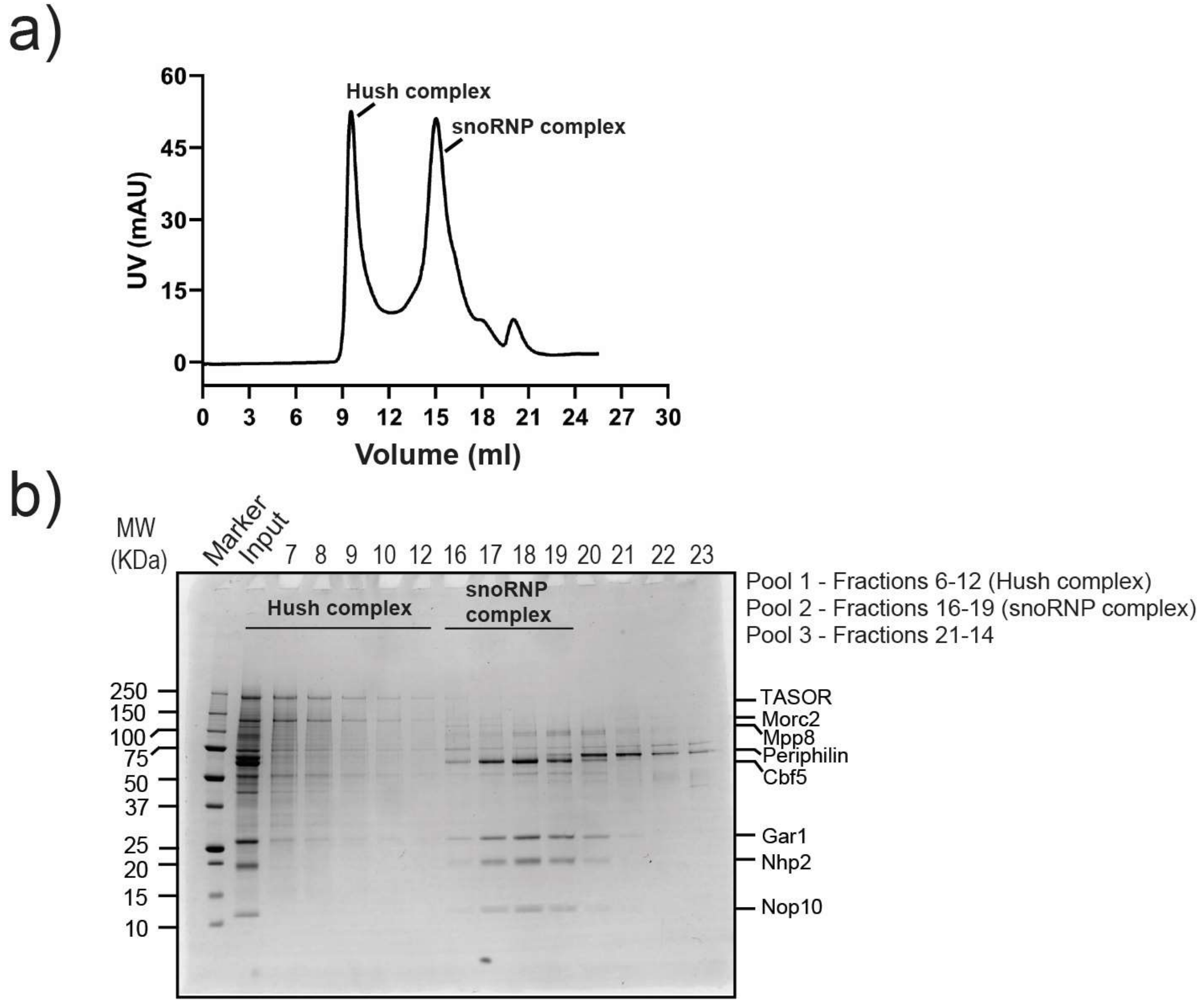


**Figure S1. Purification of Hush complex.** A) Superose 6 size exclusion chromatogram showing elution of hush complex (peak 1) and H/ACA snoRNP complex (peak 2). B) SDS-PAGE gel bands of purified HUSH and H/ACA snoRNA complex.

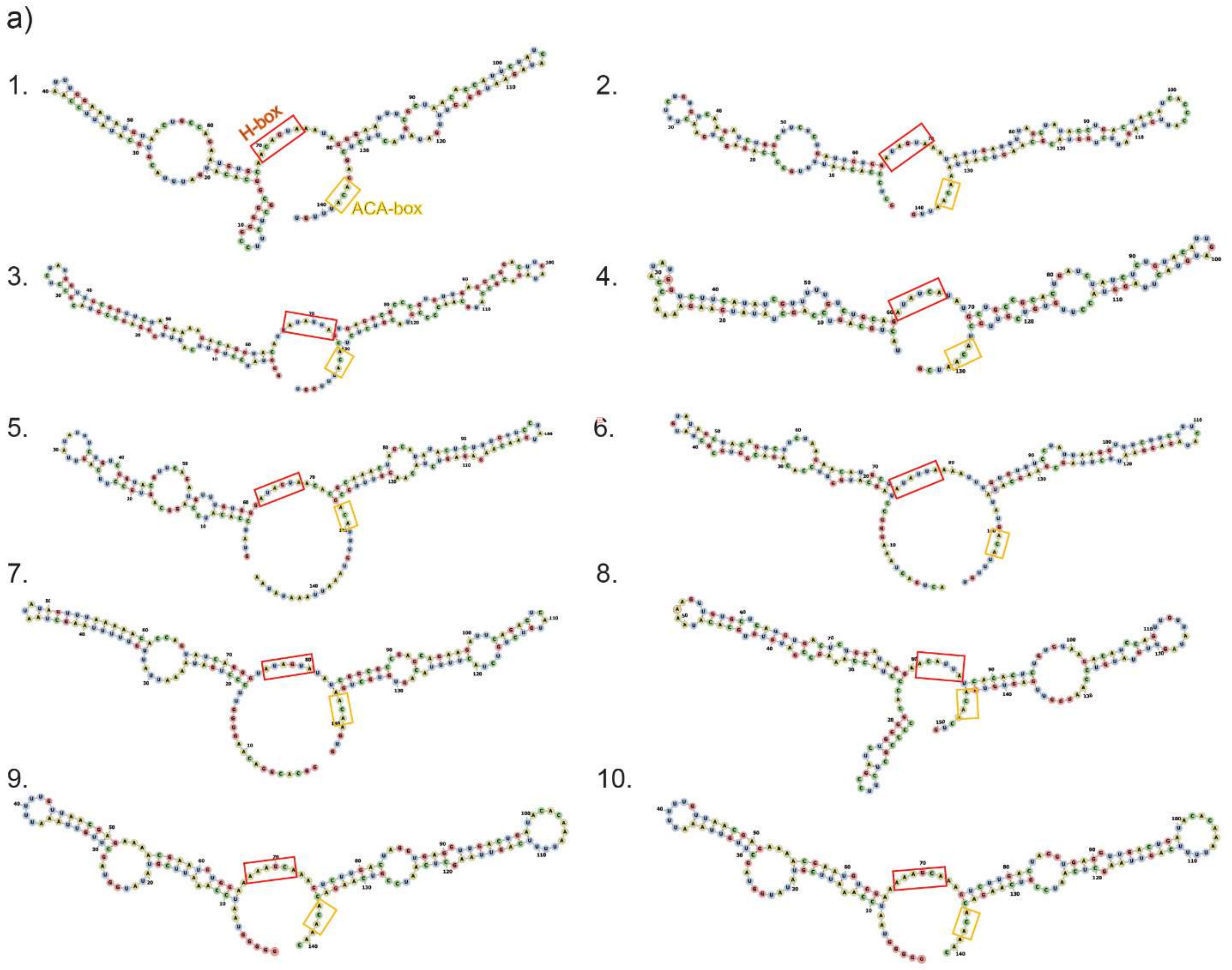


**Figure S2. The most abundant snoRNAs from the purified snoRNP complexes.** The snoRNA secondary structure prediction is done by the prediction tool - Mxfold2. The H boxes and ACA boxes are highlighted by orange and yellow boxes respectively.

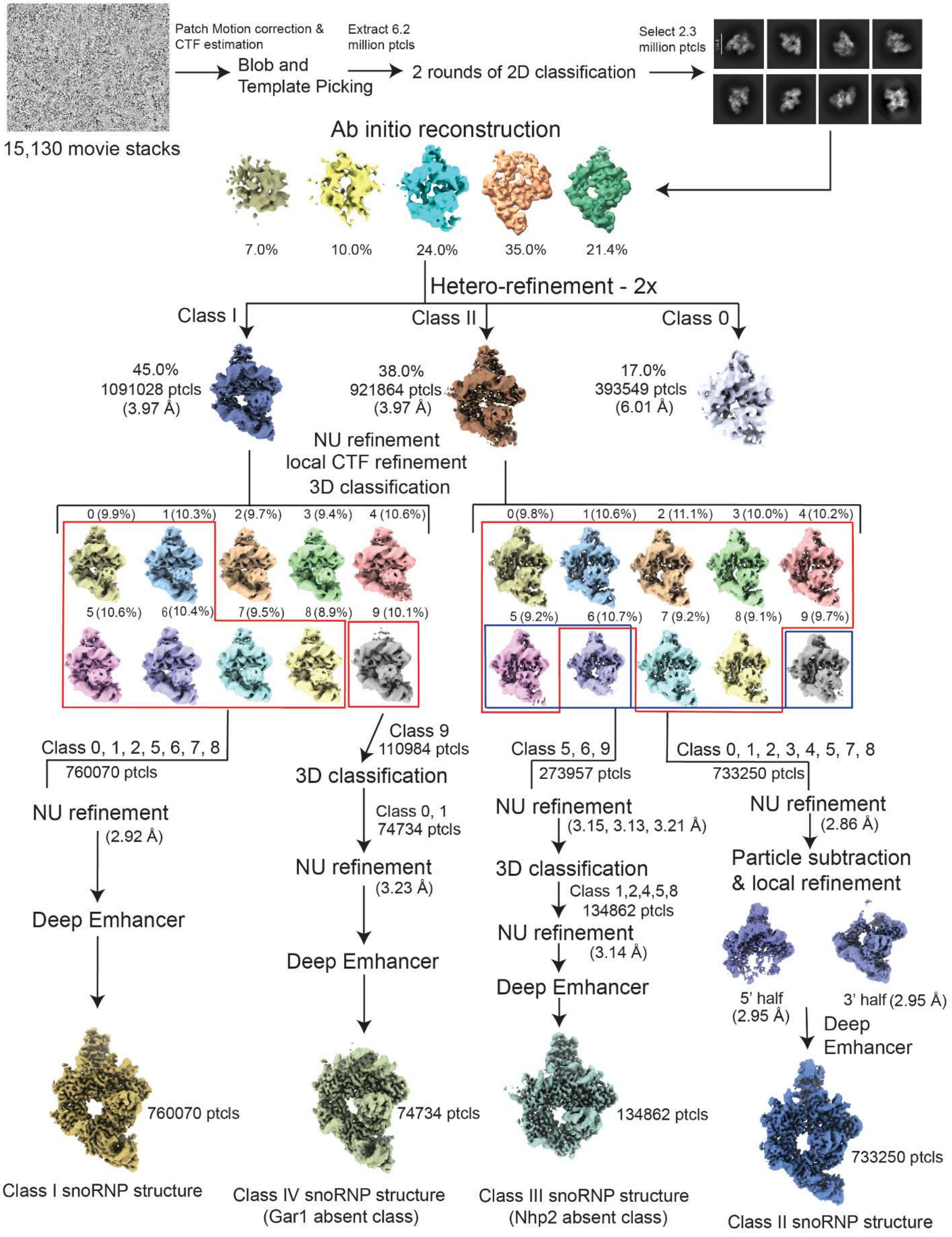


**Figure S3. Cryo-EM processing pipeline for the insect H/ACA snoRNP complex.**

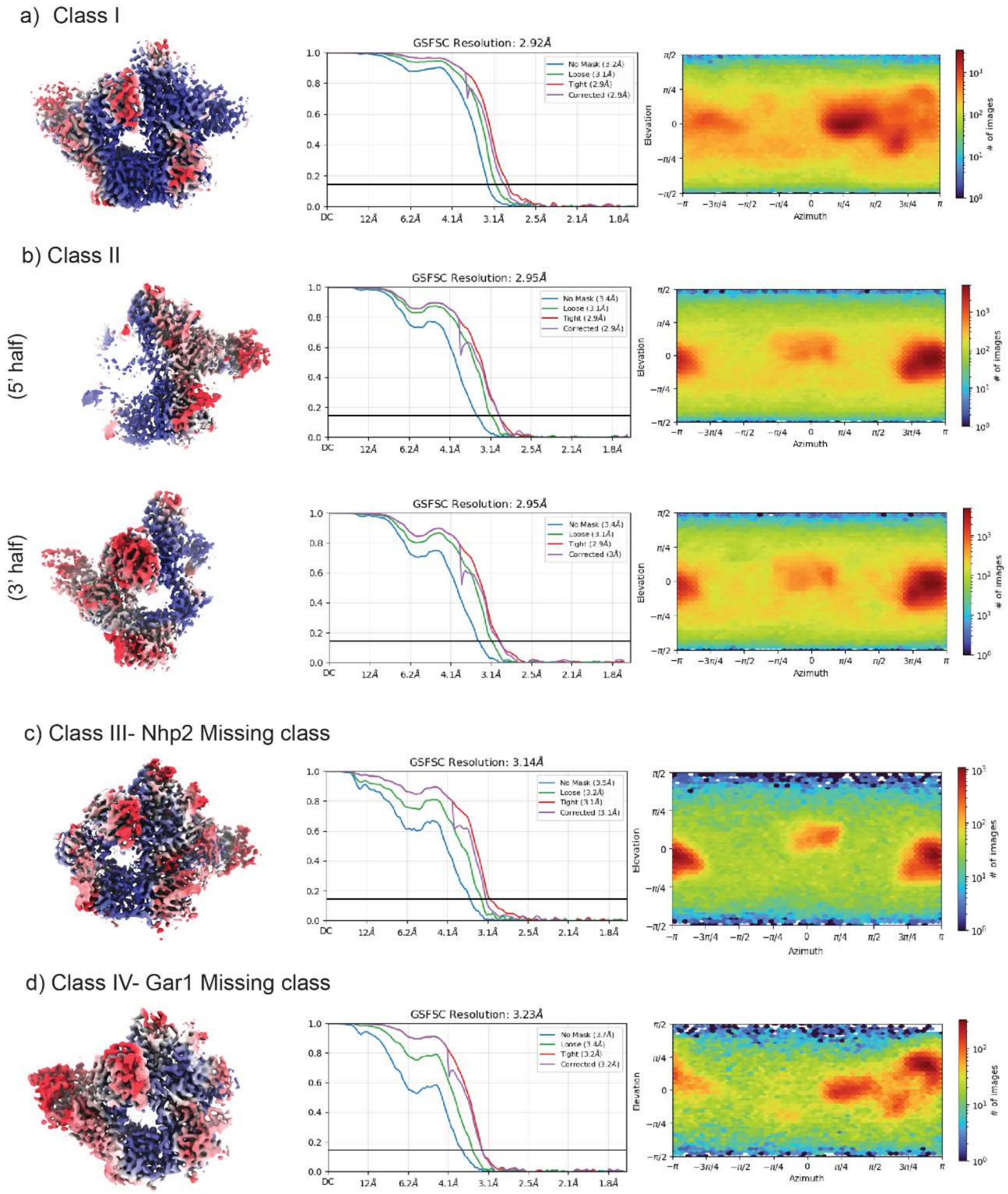


**Figure S4. Cryo-EM map validation of the four H/ACA snoRNP classes.** Local resolution estimates (left), FSC curves (middle) and particle orientation distribution (right) for, **a)** class I, **b)** class II, **c)** Nhp2 absent class III and **d)** Gar1 absent class IV.

a)

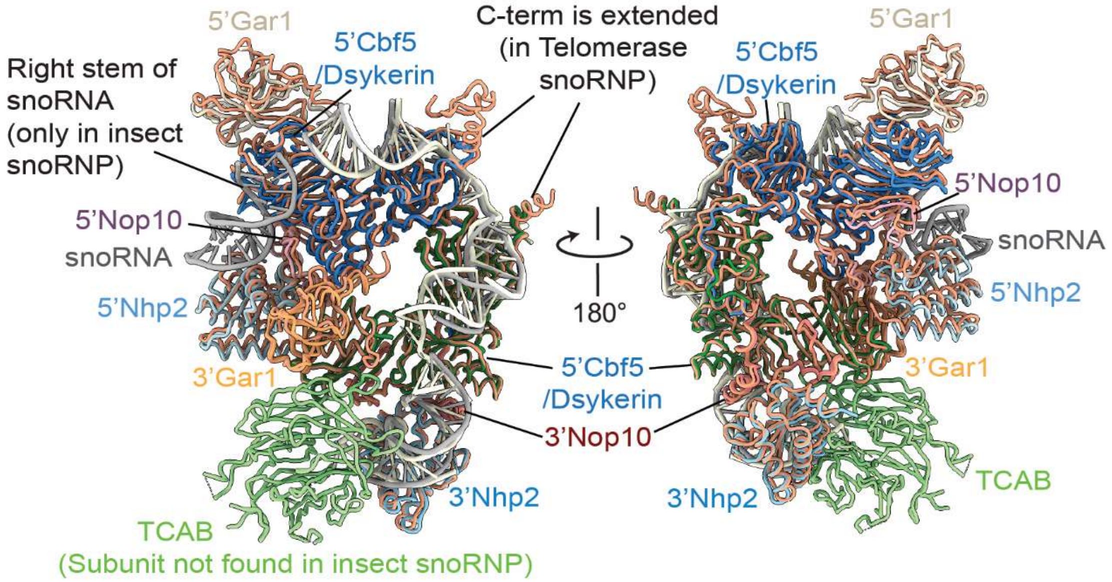


**Figure S5. Comparison between *T.ni* H/ACA snoRNP and human telomerase complex.** The insect H/ACA snoRNP structure (colored as in Fig.1) superimposed with the human telomerase H/ACA module (PDB ID – 8OUF, shown in light salmon, with TCAB shown in light green). Differences between the two structures are highlighted.

### Sequence Alignment (With Dsykeratosis Congenita mutations/substitutions marked with * )

#### a) Dyskerin/Cbf5

```
                                                                     *****  *     *
Conservation:               9                                  6    9 6 6        966 9999999696966
DKC1_T.ni                1  MTEVLQPGAEVFS--EKKKKKVKESK-EGVSLGQFQKLGDFKIEPTE-SVTKLDTAYWPLLLKNFDRLNV   66
Cbf5_S.Cerevisiae        1  M-----------------------------------SKEDFVIKPEA-AGASTDTSEWPLLLKNFDKLLV   34
DKC1_H.sapiens           1  MAD-----AEVIILPKKHKKKKERKSLPEEDVAEIQHAEEFLIKPES-KVAKLDTSQWPLLLKNFDKLNV   64
DKC1_D.melanogaster      1  MAD-----VEVRK--EKKKKKIKEEPLDGDDIGTLQKQGNFQIKPSS-KIAELDTSQWPLLLKNFDKLNI   62
DKC1_M.musculus          1  MAD-----AEVITFPKKHKKKKDRKPLQEDDVAEIQHAEEFLIKPES-KVAQLDTSQWPLLLKNFDKLNV   64
Cbf5_S.pombe             1  M--------------------------------TDTHPGVDFMIKPEATSASKIDTAEWPLLLKNFDKLLV   39
DKC1_C.elegans           1  M-----------------GKKDKRSKLEGDDLAEAQQKGSFQLPSSN-ETAKLDASQWPLLLKNYDKLNV   52
Consensus_aa:               M.............................sphpp..pFbIcPps.phtplDTtpWPLLLKNFDKL.V

                            ****   *                                                *
Conservation:               96 9999   9 69969 6  9  9669699969999999969666996699999999999999699996
DKC1_T.ni               67  RTNHYTPLPFGHSPLKRPIAEYVKAGFINVDKPSNPSSHEVVSWIKRILKVEKTGHSGTLDPKVTGCLIV  136
Cbf5_S.Cerevisiae       35  RSGHYTPIPAGSSPLKRDLKSYISSGVINLDKPSNPSSHEVVAWIKRILRCEKTGHSGTLDPKVTGCLIV  104
DKC1_H.sapiens          65  RTTHYTPLACGSNPLKREIGDYIRTGFINLDKPSNPSSHEVVAWIRRILRVEKTGHSGTLDPKVTGCLIV  134
DKC1_D.melanogaster     63  RSNHYTPLAHGSSPLNRDIKEYMKTGFINLDKPSNPSSHEVVAWIKKILKVEKTGHSGTLDPKVTGCLIV  132
DKC1_M.musculus         65  RTAHYTPLPCGSNPLKREIGDYIRTGFINLDKPSNPSSHEVVAWIRRILRVEKTGHSGTLDPKVTGCLIV  134
Cbf5_S.pombe            40  RTGHYTPIPCGNNPLKRPIAEYVSSGVINLDKPANPSSHEVVAWVKKILRVEKTGHSGTLDPKVTGCLII  109
DKC1_C.elegans          53  RTNHYTPHVEGVSPLKRDIKNYISSGFFNLDKPSNPSSHEVVSWIKRILRCEKTGHSGTLDPKVSGCLIV  122
Consensus_aa:               RosHYTPlshGssPLKR.I.pYlpoGhINLDKPSNPSSHEVVtWI++IL+hEKTGHSGTLDPKVTGCLIV

                                                   *
Conservation:               9669699969999 999996 6  99   6    6   69 96999999999996999999969669666
DKC1_T.ni              137  CIDRATRLVKSQQNAGKEYVAVFSLHSAVENVKKVTQGLEKLRGALFQRPPLISAVKRQLRVRSVYDSKL  206
Cbf5_S.Cerevisiae      105  CIDRATRLVKSQQGAGKEYVCIVRLHDALKDEKDLGRSLENLTGALFQRPPLISAVKRQLRVRTIYESNL  174
DKC1_H.sapiens         135  CIERATRLVKSQQSAGKEYVGIVRLHNAIEGGTQLSRALETLTGALFQRPPLIAAVKRQLRVRTIYESKM  204
DKC1_D.melanogaster    133  CIDRATRLVKSQQSAGKEYVAIFKLHGAVESVAKVRQGLEKLRGALFQRPPLISAVKRQLRVRTVYDSKL  202
DKC1_M.musculus        135  CIERATRLVKSQQSAGKEYVGIVRLHNAIEGGTQLSRALETLTGALFQRPPLIAAVKRQLRVRTIYESKM  204
Cbf5_S.pombe           110  CNDRATRLVKSQQSAGKEYVCVLRLHDSVEGERNVASAIETLTGALFQRPPLISAVKRQLRIRSIYESKL  179
DKC1_C.elegans         123  CIDRTTRLAKSQQGAGKEYICIFKLHEEVEDDRKVKQALEKLTGALFQRPPLISAVKRQLRIRTVYENKF  192
Consensus_aa:               CI-RATRLVKSQQtAGKEYVtlh+LHstlEs.ppl.ptLEpLpGALFQRPPLItAVKRQLRlRolY-SKh

Conservation:               6669  6  6696 6996969 99 999996 9969969 99996999   9 6 699699669966 66
DKC1_T.ni              207  LDFDKDRNIGVFWVSCEAGSYIRTMCVHLGLMLGVGGQMIELRRVRSGIQGEKEGMVTMHDILDAQWAYE  276
Cbf5_S.Cerevisiae      175  IEFDNKRNLGVFWASCEAGTYMRTLCVHLGMLLGVGGHMQELRRVRSGALSENDNMVTLHDVMDAQWVYD  244
DKC1_H.sapiens         205  IEYDPERRLGIFWVSCEAGTYIRTLCVHLGLLLGVGGQMQELRRVRSGVMSEKDHMVTMHDVLDAQWLYD  274
DKC1_D.melanogaster    203  LDYDETRNMGVFWVSCEAGSYIRTMCVHLGLVLGVGGQMLELRRVRSGIQSERDGMVTMHDVLDAMWLYE  272
DKC1_M.musculus        205  IEYDPERRLGIFWVSCEAGTYIRTLCVHLGLLLGVGGQMQELRRVRSGVMSEKDHMVTMHDVLDAQWLYD  274
Cbf5_S.pombe           180  IEFDNERNLAVFWASCEAGTYMRTLCVHLGLLLGVGGHMQELRRVRSGCLSENDDIVTMHDVLDAQWIYD  249
DKC1_C.elegans         193  IEYDPAQQMGIFNCICESGTYVRTICVHLGLILGCGGQMQELRRNRSGICDENENMVTMHDVLDAQYLLD  262
Consensus_aa:               l-@D.pRphGlFWhSCEAGoYhRThCVHLGLlLGVGGpMbELRRVRSGhbtEp-.MVTMHDVLDAQWlY-

                                 *                       *         *  *    *
Conservation:               6 6999669 66 999 996 699   999 699699999  699 9996 699   999  69999966
DKC1_T.ni              277  NHKDESYLRRVIKPLEGLLVAHKRIFIKDSAVNAVCYGAKVLLPGILRYEDGIEIDQEIVIVTTKGEAVA  346
Cbf5_S.Cerevisiae      245  NTRDESYLRSIIQPLETLLVGYKRIVVKDSAVNAVCYGAKLMIPGLLRYEEGIELYDEIVLITTKGEAIA  314
DKC1_H.sapiens         275  NHKDESYLRRVVYPLEKLLTSHKRLVMKDSAVNAICYGAKIMLPGVLRYEDGIEVNQEIVVITTKGEAIC  344
DKC1_D.melanogaster    273  NHKDESMLRRVIKPLEGLLVNHKRIIMKDSSVNAVCYGAKITLPGVLRYEDGIEIDQEIVICTTKGEAIC  342
DKC1_M.musculus        275  NHKDESYLRRVVYPLEKLLTSHKRLVMKDSAVNAICYGAKIMLPGLLRYEDGIEVNQEIVVITTKGEAIC  344
Cbf5_S.pombe           250  NTRDESYLRRVIRPLESLLVGYKRIVVKDSAVNAICYGAKLMIPGLLRYEAGIEVNEEIVLITTKGEAIA  319
DKC1_C.elegans         263  TQKDESYMRHIVRPLEALLTQHKRVVVKDSCINAICYGAKILIPGILRYDDDIEVGKEIVIMSTKGEAIC  332
Consensus_aa:               Nh+DESYLR+ll.PLE.LLhs@KRllhKDStVNAlCYGAKlhlPGlLRYE-GIElspEIVlhTTKGEAIt

                                 *  *   * *                  *     * *           *         **
Conservation:                69969 9   6669996696 99699999 969699999 6  99 6   96999 96666 99  9
DKC1_T.ni              347  LAIALMTTSTMASCDHGVAAKLKRVIMERDTYPRKWGLGPKASHKKSLISQGKLDKFGKPNENTPSEWLN  416
Cbf5_S.Cerevisiae      315  VAIAQMSTVDLASCDHGVVASVKRCIMERDLYPRRWGLGPVAQKKKQMKADGKLDKYGRVNENTPEQWKK  384
DKC1_H.sapiens         345  MAIALMTTAVISTCDHGIVAKIKRVIMERDTYPRKWGLGPKASQKKLMIKQGLLDKHGKPTDSTPATWKQ  414
DKC1_D.melanogaster    343  LAIALMTTATMASCDHGVVAKIKRVIMERDTYPRKWGLGPKASAKKALIAAGKLDKFGRPNENTPKEWLT  412
DKC1_M.musculus        345  MAIALMTTAVISTCDHGIVAKIKRVIMERDTYPRKWGLGPKASQKKMMIKQGLLDKHGKPTDNTPATWKQ  414
Cbf5_S.pombe           320  VGIAQMSTVELSTCDHGVVAKVKRCIMERDVYPRRWGLGPQSMKKKTLKKEGKLDKYGRPNENTPADWSK  389
DKC1_C.elegans         333  IAIAQMNTSTIASVDHGVVAKSKRVIMERDVYGRKWGLGPVASKKKQMVKDGLLDKFGKPNDTTPKSWAK  402
Consensus_aa:               hAIAbMoTsshtoCDHGlVAKlKRhIMERDhYPR+WGLGP.AppKK.hb.pGbLDK@G+Ps-sTP.pW.p

Conservation:                96                                    6  6 6                  66
                                 *
DKC1_T.ni              417  SYVDYKK-------KDT-------------NGDAEDDVGRKRTAST------ANADDPDNSVEVKS--EK  458
Cbf5_S.Cerevisiae      385  EYVPLDN-----AEQSTSSSQETKETEEEPKKAKEDSLIKEVETEKEEVKEDDSKKEKKEKKDKKEKKEK  449
DKC1_H.sapiens         415  EYVDYSE-----SAKKEVVAEVVKAPQVVAEAAKTAKRKRESESES---DETPPAAPQLIKKEKK-----  471
DKC1_D.melanogaster    413  GYVDYNA-------KKPAAQEVSPT-----NGSSEPSKRKLSTSSV---EETAAAAVSEETPSKDK--KK  465
DKC1_M.musculus        415  DYIDYSD-----SGKNTLVTEAVQAPQLAAEAVNVIKRKRDSESES---DET-PTVPQL--KEKK-----  468
Cbf5_S.pombe           390  SYIDYSDPNAEVAKPAPVVAPAAPTVEAEVNGVEDSKKRKSVESSE---KDEDEAAKKEEKRRKKE--AK  454
DKC1_C.elegans         403  EYVQTST-------KKEVKKEETPDEE-----------------------------EEEAPKKKS--KK  433
Consensus_aa:               pYlshsp.......pp.h..p.h.s.....ptsppsp..+pspopp.......shs.pbpp.cKK.....

Conservation:               6 96  6                                  6 9969969
DKC1_T.ni              459  KKKKKKRDSEAADAVPEDGSGDATMEAETTTEADSSIRKEKKKKKKKDREQER--VDE-----  514
Cbf5_S.Cerevisiae      450  KEKKDKKE------------------------------KKEKKEKKRKSEDGDSEEKKSKKSKK  483
DKC1_H.sapiens         472  KSKKDKKAKAGLESGAEPGDGD-------------SDTTKKKKKKKKAKEVEL--VSE-----  514
DKC1_D.melanogaster    466  KKKKHKGDEEAPEAAEEEAE---------------PVEKEKKKKKKKDKDRDRDEAQE-----  508
DKC1_M.musculus        469  -KKKDKKPKTVLESGGETGDGD-------------NDTT-KKKKKKVKVVEE--MSE-----  509
Cbf5_S.pombe           455  KEKKEKKE------------------------------KKEKKEKKKKSE-------------  474
DKC1_C.elegans         434  SKKQESSDSDSD--------------------------------------------------  445
Consensus_aa:               KcKKcKp..........................pp.KKcKKKKsc.............
```

**b) Gar1**

```
Conservation:             969                            99        6  699 69  6696669  99696
Gar1_T.ni              1  MSF-RGRGGGGGGRGGYGGGGGG-GG-----RGF----GGRG-GGGGRGGFGGRGGGRG--GRGGGGGGF   56
Gar1_S.cerevisiae      1  MSF----------------------------RG--------G-NRGGRGG--FRGGFRG--GRTGSARSF   29
Gar1_H.sapiens         1  MSF-RGGGRGGFNRGGGGGGFNR-GGSSNHFRG--------G--GGGGGGGNFRGGGRGGFGRGGGRGGF   58
Gar1_D.melanogaster    1  MGFGKPRGGGGGGGRGFGGGGGG-GG-----RGFGGGGGGRG-GGGGRGGGGGFGRGGG--GRGGGRGAF   61
Gar1_M.musculus        1  MSF-RGGGRGGFNRGGGGGGFNRGGGSNNHFRGGGGGGGGSFRGGGGGGGGSFRGGGRGGFGRGGGRGGF   69
Gar1_S.pombe_          1  MSF----------------------------RG--------G--RG--------GGFRG--GRGGSRPFT   22
Gar1_C.elegans         1  MSFRGGRGGGGGFRGGRGGGGGG-GF-----RG--------G--RGGDRGGGFR-GGRGGFG-GGGRGGY   52
Consensus_aa:             MSF............................RG........G...GG.GG...RGG.RG..GRGGt.st@

Conservation:                   9996 9   9 6   66 6 69        6999999666699 9699699669      6666
Gar1_T.ni             57  RQ-QDQGPPESVIPLGHYGWTVQDDLICKVDIED--VPYFNAPIFLENKEQIGKIDEIFGNLRDYFVSVK  123
Gar1_S.cerevisiae     30  ----QQGPPDTVLEMGAFLHPCEGDIVCRSINTK--IPYFNAPIYLENKTQVGKVDEILGPLNEVFFTIK   93
Gar1_H.sapiens        59  NKGQDQGPPERVVLLGEFLHPCEDDIVCKCTTDENKVPYFNAPVYLENKEQIGKVDEIFGQLRDFYFSVK  128
Gar1_D.melanogaster   62  ----DTGPPERVIPLGNYVYSCQNDLVCKVDIQD--VPYFNAPIFLENKEQVGKIDEIFGTVRDYSVSIK  125
Gar1_M.musculus       70  NKFQDQGPPERVVLLGEFMHPCEDDIVCKCTTEENKVPYFNAPVYLENKEQVGKVDEIFGQLRDFYFSVK  139
Gar1_S.pombe_         23  ----PSGPPDQVIELGLFMHDCEGEMVCQSTNVK--IPYFNAPIYLENKSQIGKIDEVFGPMNQVYFTVK   86
Gar1_C.elegans        53  ----DQGPPEEVVLVGVFSHQCQDDIVCNNTSGK--IPYFNAPIYFKNKEQVGKIDEIFGSPGENGFSVT  116
Consensus_aa:             ....spGPP-pVl.lG.@.@sCpsDlVCpss..c..lPYFNAPl@LENKpQlGKlDEIFG.hp-h.holK

Conservation:              6 6  6 996   6 66   9969  9999 6   6   6           69699  6999   9699
Gar1_T.ni            124  MGDNFKANSFKDGQQFYIDPAKLLPLKRFLPQPP--GAKRGGGRGRGGPRGGRGGGGGF-GRG---G--G  185
Gar1_S.cerevisiae     94  CGDGVQATSFKEGDKFYIAADKLLPIERFLPKPKVVGPP-KPKNKKKRSGAPGGRGGASMGRG---GSRG  159
Gar1_H.sapiens       129  LSENMKASSFKKLQKFYIDPYKLLPLQRFLPRPP--GEK-GPPR-----------GGGRGGRG---GGRG  181
Gar1_D.melanogaster  126  LSDNVYANSFKPNQKLFIDPGKLLPIARFLPKPP--QPK-GAKKAFTNNRGGGGGGGGFGGRG---GGRG  189
Gar1_M.musculus      140  LSENMKASSFKKLQKFYIDPYKLLPLQRFLPRPP--GEK-GPPR--------GGGGGGRGGRG---GGRG  195
Gar1_S.pombe_         87  PSEGIVSSSFKVGDKVYLSGDKLIPLDRFLPKPKTVGPK-KPKGARNGPAGRGGRGGFRGGRG---GSRG  152
Gar1_C.elegans       117  LSQGVKASSFEPESQLYIDPGKLLPVDRFLPQAG--GGR-G--------RGGRGRGGDRGGRGSDRGGRG  175
Consensus_aa:             ht-shbAsSFK..pphYIss.KLLPlpRFLPpP...G.+..s..........G.GGs.GGRG...GtRG

Conservation:             96666 6   66  69 999           69  6996669 69  96696
Gar1_T.ni            186  GGGRGGGFGRGG--GGGRGG--FGRGGGGGYGGG-GGGFNRGGGGFRGGRG-GR----------------  233
Gar1_S.cerevisiae    160  GFRGGRG---GSSFRGGRGGSSF---RGGSRGGSFRGGSRGGSRG--GFRGGRR----------------  205
Gar1_H.sapiens       182  GGGRGGG---RG--GGFRGG-----R-----GGG-GGGFRGGRGG--GFRGRGH----------------  217
Gar1_D.melanogaster  190  GGGRGGGGGRGG--GGFRGG--AGRNGG---GGG-GGGFNRGRGGG-GGGGGGRGRW-------------  237
Gar1_M.musculus      196  GGGRGGG---RG--GGFRGG-----------RGG-GGGFRGGRGGG-GFRGRGH----------------  231
Gar1_S.pombe_        153  GFGGNSR---GGFGGGSRGG--F---GGGSRGGS-RGGFRGGSRG--GFRGRF-----------------  194
Gar1_C.elegans       176  GFGRGGG---GGFRGGDRGG--FGGGRGGFRGGD-RGGFRGGRGGDFGGRGRGDFKRSYDGGSFGGQNNK  239
Consensus_aa:             G.G.GtG....G..GG.RGG..........GGt..GGFp.Gp.G..G.RG..+................

Conservation:
Gar1_T.ni                 -----
Gar1_S.cerevisiae         -----
Gar1_H.sapiens            -----
Gar1_D.melanogaster       -----
Gar1_M.musculus           -----
Gar1_S.pombe_             -----
Gar1_C.elegans       240  RTKFE  244
Consensus_aa:             .....
```

**c) Nop10**

```
                                                             *
Conservation:               9 9 9 6   5 9 9996   5 9  9 569999999599595 9 559999 56 69 5
Nop10_T.ni               1  MYLRYYLNENGDRQYTLATIDPYGKPTISAHPARFSPEDKYSRHRIIIKKRFGLLLTQQPEPIL   64
Nop10_S.cerevisiae       1  MHLMYTLGPDGKRIYTLKKVTESGEITKSAHPARFSPDDKYSRQRVTLKKRFGLVPGQ------   58
Nop10_H.sapiens_         1  MFLQYYLNEQGDRVYTLKKFDPMGQQTCSAHPARFSPDDKYSRHRITIKKRFKVLMTQQPRPVL   64
Nop10_D.melanogaster     1  MYLMYTINENGDRVYTLKKRTEDGRPTLSAHPARFSPEDKYSRQRLTIKKRFGLLLTQKPEPIY   64
Nop10_M.musculus         1  MFLQYYLNEQGDRVYTLKKFDPMGQQTCSAHPARFSPDDKYSRHRITIKKRFKVLMTQQPRPVL   64
Nop10_S.pombe            1  MHLMYYLNDEGKRVYTLKKVSPDGRVTKSSHPARFSPDDKYSRQRYTLKKRFHVLLTQLPAKPY   64
Nop10_C.elegans_         1  MFLRYFLDENQQRVYTLKRTAPSGEQTLTAHPARFSPEDKNSKYRIIIKKRFGLLPTQKAKTVC   64
Consensus_aa:               M@L.YhLs-pGcRlYTLK+hs..Gp.T.SAHPARFSP-DKYSRpRlhlKKRF.lL.TQbs...h
```

**d) Nhp2**

```
Conservation:                  9 9 6                        9          696969 669 69   9 6996
Nhp2_T.ni               1    MGKIKQEPVEQEDQADVS------VKNEPQSYDEKVDHCSVIAKPMAPKKLSKKIYKLIKKSTSHKNYIR    64
Nhp2_S.cerevisiae       1    MGKDNKEHKESK------------ESKTVDNYEARMPAVLPFAKPLASKKLNKKVLKTVKKASKA-KNVK    57
Nhp2_H.sapiens          1    MTKIKADPDGPEAQAE--------ACSGERTYQELLVNQNPIAQPLASRRLTRKLYKCIKKAVKQ-KQIR    61
Nhp2_D.melanogaster     1    MGKVKVERSEDADESVVASGD--VTIKEEESYDDKLIFVNAIAKPMAGKKLAKKCYKLVKKAMKHKTFLR    68
Nhp2_M.musculus         1    MTKVKAAPEESEAQAE--------GCSEERTYKELLVNLNPIAQPLASRRLTRKLYKCIKKAVKQ-KQIR    61
Nhp2_S.pombe_           1    MAKDKKDHKHS--------------GSTEDEYDSYLPALMPIAKPLAPKKLNKKMMKTVKKASKQ-KHIL    55
Nhp2_C.elegans          1    MGKRNLDETMNESTVSEANGDATAPTTEKDEYQALCELVNPIAQPLANRKLAKKVYKLIKKASAGDKTLR    70
Consensus_aa:                MsK.p.-.p.sc............s.pp.cpYp.bh..h.sIApPhAs++Ls+KhhKhlKKAsp..p.l+

                                                                                     *
Conservation:                 9 9 966 66 69696   996  9 66 6969  996   99 66696  69 9    96    6
Nhp2_T.ni              65    NGLKIVQKQLRLGEKGIVFFAGDISPIEIMCHLPAVCEEKDIPYCYTPSRKDIGAAMGTMRGCVMVLVKE   134
Nhp2_S.cerevisiae      58    RGVKEVVKALRKGEKGLVVIAGDISPADVISHIPVLCEDHSVPYIFIPSKQDLGAAGATKRPTSVVFIVP   127
Nhp2_H.sapiens         62    RGVKEVQKFVNKGEKGIMVLAGDTLPIEVYCHLPVMCEDRNLPYVYIPSKTDLGAAAGSKRPTCVIMVKP   131
Nhp2_D.melanogaster    69    NGLKDVQTRLRKGETGICIFAGDVTPVDIMCHLPAVCEEKGIPYTYTPSRADLGAAMGVKRGTVALLVRQ   138
Nhp2_M.musculus        62    RGVKEVQKFVNKGEKGIMVLAGDTLPIEVYCHLPVLCEDQNLPYVYIPSKTDLGAATGSKRPTCVIMVKP   131
Nhp2_S.pombe_          56    RGVKEVVKAVRKGEKGLVILAGDISPMDVISHIPVLCEDNNVPYLYTVSKELLGEASNTKRPTSCVMIVP   125
Nhp2_C.elegans         71    EGIKDVQKELRRNEKGICILAGNVSPIDVYSHIPGICEEKEIPYVYIPSREQLGLAVGHRRPSILIFVKP   140
Consensus_aa:                pGlK-V.K.lp+GEKGlhlhAGDh.Ph-lhtHlPhlCE-pslPYhYhPS+ppLGhA.ts+Rsoshlhl..

                                           *
Conservation:                          6666 6 6   6   9
Nhp2_T.ni             135    HD-----------DYKDLFDEVRGEIKLLGHPI-   156
Nhp2_S.cerevisiae     128    GSNKKKDGKNKEEEYKESFNEVVKEVQAL-----   156
Nhp2_H.sapiens        132    HE-----------EYQEAYDECLEEVQSLPLPL-   153
Nhp2_D.melanogaster   139    NE-----------EYKDLYDEVKEELSALNIPV-   160
Nhp2_M.musculus       132    HE-----------EYQETYDKCLEEVQALPTPL-   153
Nhp2_S.pombe_         126    GGKKKD--MSKVEEYKESYEEIIKEVPALEV---   154
Nhp2_C.elegans        141    SG-----------DFKELYDEVAEALRHLTVEAA   163
Consensus_aa:                .............-Yp-.@-Eh.cElphLsh...
```

**Figure S6. Sequence Alignment of snoRNA proteins protein sequences from eukaryotes.** Sequences of Dkc1/Cbf5, Gar1, Nop10 and Nhp2 (A-D) was aligned from different eukaryotic species – *Trichoplusia ni*, Saccharomyces cerevisiae, Homo sapiens, Drosophila melanogaster, Mus musculus, Saccharomyces pombe, C. elegans prepared using PROMAL3SD. The dyskeratosis congenita mutations/substitutions are marked with a star. Also, shown is the conservation of residues across the species (on the top, 9 represent the highest conservation index). Consensus amino acids are indicated at the bottom of the alignment. Representative sequences have magenta names, and they are colored according to predicted secondary structures (red: alpha-helix, blue: beta-strand).

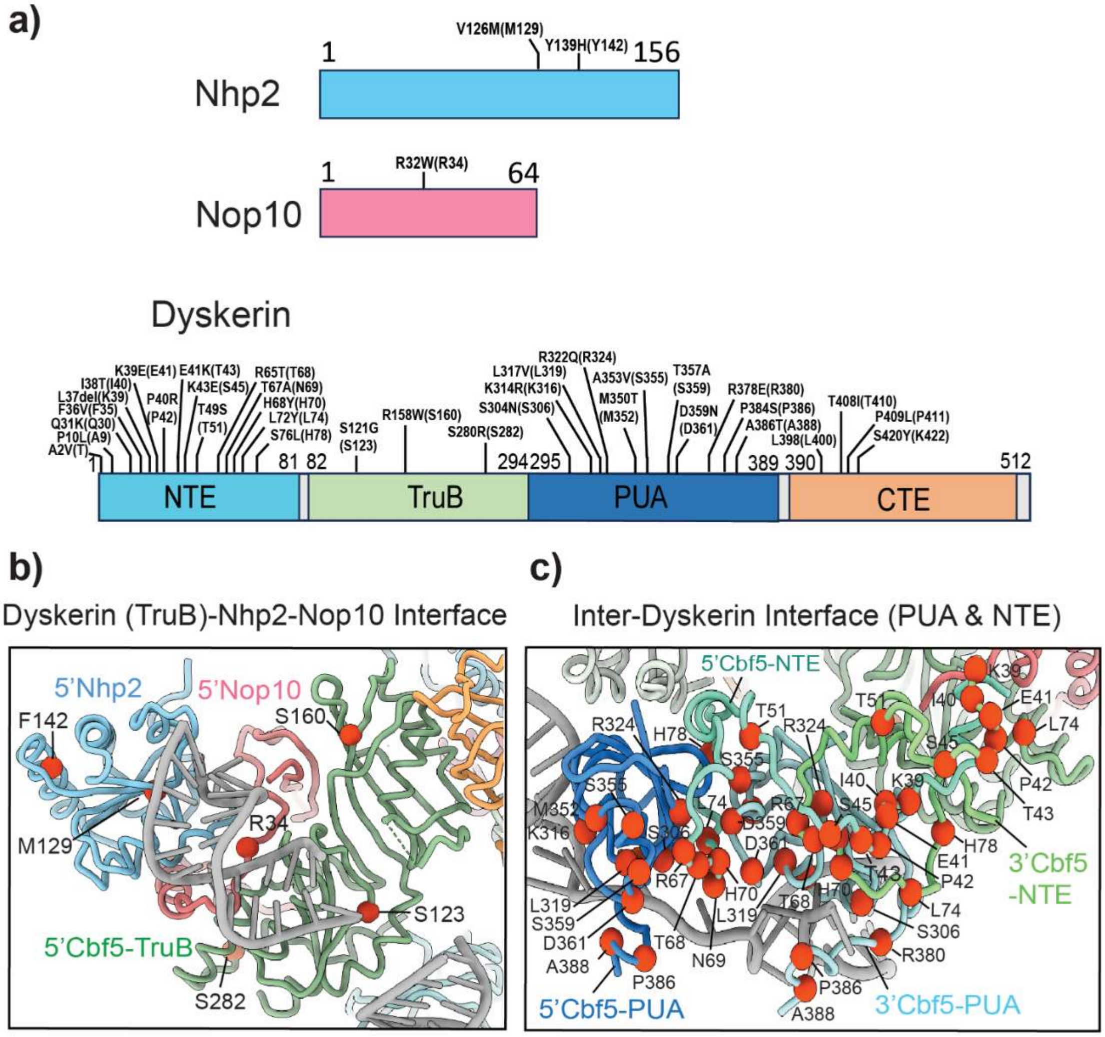


**Figure S7. Dyskeratosis congenita mutations (DC) are mapped on the insect H/ACA snoRNP structure. a)** Schematic representation of Cbf5, Nhp2, Nop10 with DC mutations/substitutions as marked in Fig.3a. **b)** Mutations highlighted for 5'Nhp2, 5'Nop10 and the TruB domain of Cbf5. **c)** Mutations highlighted for the PUA and NTE domains of Cbf5. Red spheres denote mutations.

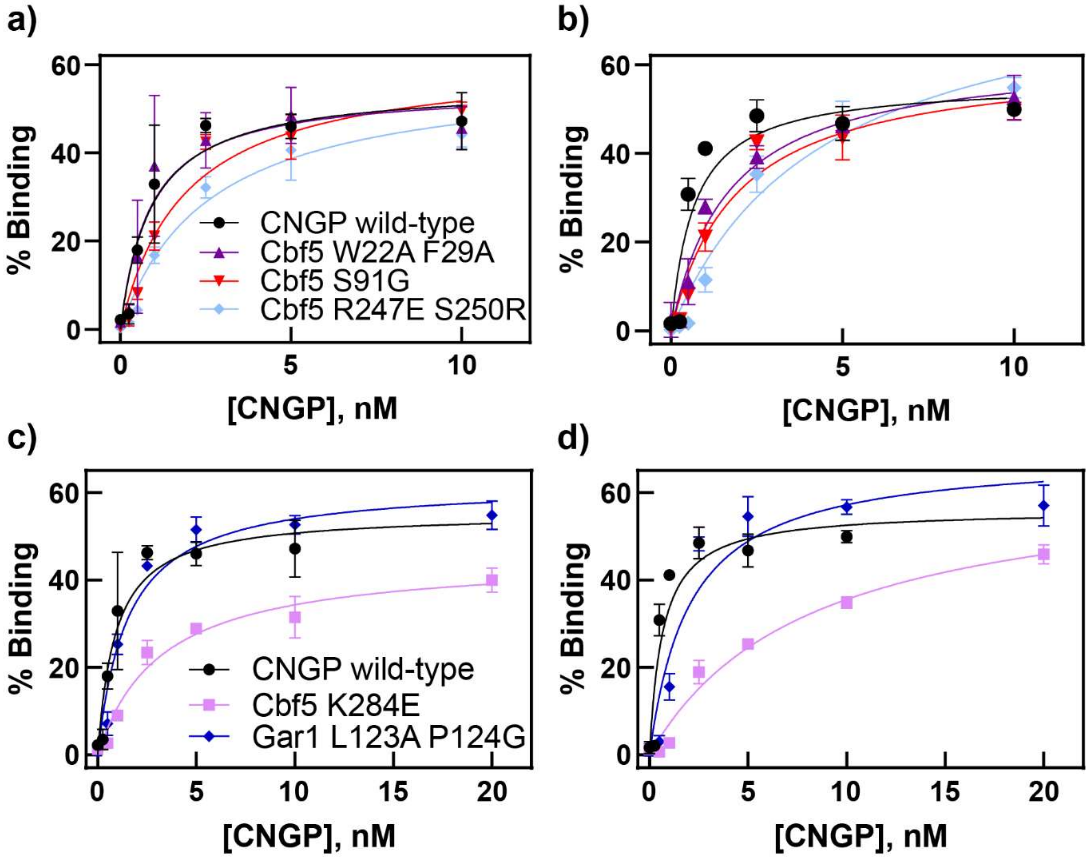


**Figure S8. Nitrocellulose filtration assays to determine binding of Cbf5-Nop10-Gar1-Nhp2 to individual hairpins of the H/ACA snoRNP snR34.** The Cbf5-Nop10-Gar1-Nhp2 (CNGP) complex was recruited and titrated with radioactively labelled 5′ hairpin (A and C, left) or 3′ hairpin (B and D, right) of snR34. The binding analysis was conducted for different protein variants: Cbf5 W22A F29A, Cbf5 S91G, Cbf5 R237E S250R **(a-b)** as well as Cbf5 K284E and Gar1 L123A P124G **(c-d)**.

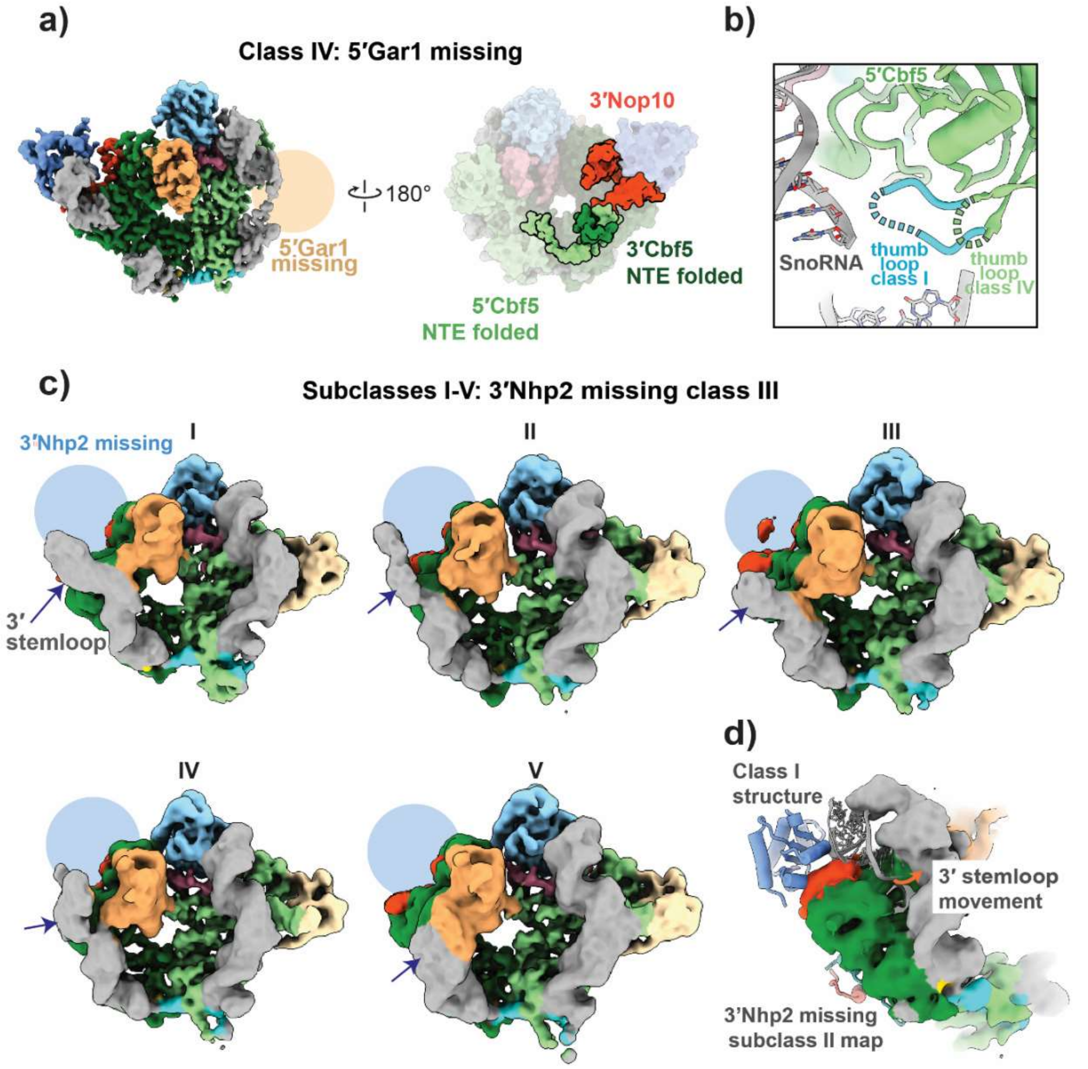


**Figure S9. Structure of the endogenous H/ACA snoRNP classes missing Gar1. (a)** Cryo-EM structure of 5′ class IV. Cryo-EM density for the structure is shown on the left. A surface representation of the refined model is shown on the right. The NTE of 5′ and 3′ Cbf5 and 3’ Nop10 are well folded. **(b)** Comparison of the thumb loop conformations in Gar1 class IV and class I. **(c)** Subclassification of class III which lacks NHP2 (subclasses I-V). The 3’ stemloop for each subclass is indicated by an arrow. The position of the Nhp2 binding site is indicated with a light blue circle. **(d)** overlay of the class I snoRNP structure with the cryo-EM map of subclass II of class III, indicating that the 3’ stemloop is moved in the subclass as compared to snoRNA when NHP2 is present.

### 1.3 Tables

### Table S1, S2, S3, S4 are attached separately

### Table S5. Cryo-EM data collection, refinement and validation statistics

**Data statistics**

| Sample | Cross-linked snoRNA | Cross-linked snoRNA | Cross-linked snoRNA | Cross-linked snoRNA |
|---|---|---|---|---|
| Microscope | Titan Krios | Titan Krios | Titan Krios | Titan Krios |
| Voltage (kV) | 300 | 300 | 300 | 300 |
| Electron exposure ($e^{-}$/Å$^2$) | 60 | 60 | 60 | 60 |
| Pixel size (Å) | 0.414 | 0.414 | 0.414 | 0.414 |
| Symmetry imposed | C1 | C1 | C1 | C1 |
| Initial particle projections | 7,128,691 | 7,128,691 | 7,128,691 | 7,128,691 |
| Final particle projections | 760,070 | 733,250 | 134,682 | 74,734 |
| Map resolution (Å) | 2.92 | 2.95 | 3.14 | 3.23 |
| FSC threshold | 0.143 | 0.143 | 0.143 | 0.143 |
| Map resolution range (Å) | 2.5 - 7.0 | 2.5 - 7.0 | 2.5 - 7.0 | 2.5 - 7.0 |

| EMD code | EMD-XXXXX | EMD-XXXX | EMD-XXXX | EMD-XXXXX |
|---|---|---|---|---|
| Structure model<br>PDB code | Class I<br>XXXX | Class II<br>XXXX | Class III<br>XXXX | Class IV<br>XXXX |
| Model resolution (Å)<br>FSC threshold | 2.92<br>0.143 | 2.95<br>0.143 | 3.14<br>0.143 | 3.23<br>0.143 |
| Model composition | | | | |
| Non-hydrogen atoms | 12320 | 11840 | 10251 | 11363 |
| Protein residues | 1272 | 1225 | 1080 | 1167 |
| RNA base pairs | 101 | 96 | 76 | 97 |
| Ligands | 0 | 0 | 0 | 0 |
| Zn ions | 0 | 0 | 0 | 0 |
| *B* factors (Å$^2$) | | | | |
| Protein | 67.63 | 81.25 | 86.91 | 64.89 |
| RNA | 109.20 | 121.24 | 124.20 | 106.84 |
| Ligands and Zn | --- | --- | --- | --- |
| R.m.s. deviations | | | | |
| Bond lengths (Å) | 0.004 (0) | 0.004 (0) | 0.004 (0) | 0.007 (0) |
| Bond angles (°) | 0.921 (0) | 0.620 (0) | 0.708 (0) | 0.674 (5) |
| Validation | | | | |
| MolProbity score | 0.99 | 1.54 | 1.36 | 1.99 |
| Clashscore | 0.71 | 4.88 | 3.37 | 7.19 |
| Rotamer outliers (%) | 0.18 | 1.02 | 0.53 | 1.86 |
| Ramachandran plot | | | | |
| Favored (%) | 96.17 | 95.93 | 96.42 | 94.26 |
| Allowed (%) | 3.82 | 4.07 | 3.58 | 5.74 |
| Disallowed (%) | 0.00 | 0.00 | 0.00 | 0.00 |
| CC (mask) | 0.78 | 0.67 | 0.63 | 0.75 |

**Table S6. Comparison of residues in insect, yeast and human H/ACA proteins.** Together with the protein, the domain location of the residue and (if relevant) the interaction with other H/ACA proteins is stated. Residues that are altered in Dyskeratosis congenita patients are indicated together with the disease substitution. Similarly, residues that were altered in yeast for functional analysis are listed including the substitution.

| **Protein (domain)** | ***Trichoplusia ni* (insect)** | ***S. cerevisiae* (baker's yeast)** | ***H. sapiens* (human)** |
|---|---|---|---|
| **5′Cbf5/Dyskerin (NTE):** At interface with 3′Cbf5 PUA | H70 | H38A | H68Q |
| | Y71 | Y39A | Y69 |
| **3′Cbf5/Dyskerin (PUA):** At interface with 5′Cbf5 NTE | M352 | M320A | M350T/I |
| | S359 | S327A | T357 |
| | D361 | D329A | D359N |
| **Cbf5/Dyskerin (TruB):** At interface with Gar1 | K279 | R247A | K277 |
| | S282 | S250R | S280R |
| **Cbf5/Dyskerin (PUA):** At interface with H/ACA snoRNA | K316 | K284E | K314R |
| **Cbf5/Dyskerin (TruB):** Close to catalytic site | S123 | S91G | S121G |
| **5′Cbf5/Dyskerin (NTE):** Interface with 3′Cbf5/Dyskerin (NTE) | W54 | W22A | W52 |
| | F61 | F29A | F59 |
| **3′Cbf5/Dyskerin (TruB):** At interface with 5′Nop10 & 5′Nhp2 | E176 | E144R | E174 |
| **Nop 10** | R51 | R51 | R43 |
| **3′Gar1:** at interface with 5′Cbf5(TruB) | L153 | L123A | L158 |
| | P154 | P124G | P159 |
| **Nhp2:** at interface with H/ACA snoRNA | K68 | K61E | K65 |
| | K72 | K65E | K70 |
| **Nhp2:** at interface Cbf5 & Nop10 | E102 | E95 | E99 |
| **Nhp2:** at interface with H/ACA snoRNA | R125 | R118E | R122 |

**Table S7. Initial velocities for pseudouridine formation in two different substrate RNAs recognized by the 5' and 3' hairpin.** Yeast H/ACA snoRNPs were reconstituted using the snR34 H/ACA snoRNA, and pseudouridylation was measured over time by tritium release (Fig. 2, 3, 4).

| **Cbf5-Nop10-Gar1-Nhp2 Variant** | **Initial velocity (nM min$^{-1}$) for 5′ substrate** | **Initial velocity (nM min$^{-1}$) for 3′ substrate** |
|---|---|---|
| wild-type | 17 ± 3.9 | 9 ± 2.6 |
| Cbf5 R247A S250R | 2 ± 3.0 | 2 ± 3.0 |
| Cbf5 W22A F29A | 10 ± 5.3 | 2 ± 2.4 |
| Cbf5 S91G | 11 ± 2.7 | 6 ± 3.9 |
| Cbf5 K284E | 8 ± 1.8 | 5 ± 2.1 |
| Gar1 L123A P124G | 6 ± 9.7 | 7 ± 3.4 |
| Nhp2 R118E | 12 ± 2.5 | 5 ± 1.5 |
| Nhp2 K61E K65E | 6 ± 2.8 | 2 ± 5.6 |
| No Nhp2 | 4 ± 1.7 | 1 ± 0.9 |

**Table S8. Dissociation constants (nM) for the binding of yeast Cbf5-Gar1-Nop10-Nhp2 variants to snR34 guide RNA.** RNA binding to the listed proteins was measured by filter binding (Fig 2, 3, 4, and Fig. S8).

| **Cbf5-Gar1-Nop10-Nhp2 Variant** | **$K_D$ (nM) for snR34 full-length** | **$K_D$ (nM) for snR34 5' hairpin** | **$K_D$ (nM) for snR34 3' hairpin** |
|---|---|---|---|
| wild-type | 1.1 ± 0.15 | 0.9 ± 0.26 | 0.7 ± 0.19 |
| Cbf5 R247A S250R | 5.2 ± 1.46 | 2.6 ± 0.53 | 4.4 ± 1.01 |
| Cbf5 W22A F29A | 1.2 ± 0.21 | 0.9 ± 0.30 | 1.7 ± 0.31 |
| Cbf5 S91G | 2.2 ± 0.37 | 1.9 ± 0.37 | 1.9 ± 0.37 |
| Cbf5 K284E | 4.6 ± 0.81 | 3.2 ± 0.56 | 8.1 ± 1.56 |
| Gar1 L123A P124G | 2.7 ± 0.41 | 1.5 ± 0.24 | 2.1 ± 0.50 |

**Table S9. Oligonucleotides used to generate yeast Nhp2 variants through site-directed mutagenesis**

| Name | Sequence (5′-3′) |
|---|---|
| Nhp2 6165 F | GTCGAGGAAGTTGTCGAGGCCTTAAGAAAGGGTGAAAAAGGTTTAGTCG TCATCGCCGG |
| Nhp2 6165 R | GGCCTCGACAACTTCCTCGACACCTCTTTTAACATTCTTGGCCTTGG |
| Nhp2 118 F | GCTACAAAAGAACCTACCTCAGTTGTCTTTATCGTCCCAGGTAGC |
| Nhp2 118 R | CTGAGGTAGGTTCTTTTGTAGCGCCAGCGGCACCTAAGTCTTGC |

**Table S10. Oligonucleotides to in vitro transcribe substrate RNAs**

| Name | Sequence (5′-3′) |
|---|---|
| H89 (5´ substrate) sense | GCTAATACGACTCACTATAGGGATAACTGGCTTGTGGCAGTCAAGCGT TCATAGCGAC |
| H89 (5´ substrate) anti-sense | CATCGAAGAATCAAAAAGCAATGTCGCTATGAACGCTTGACTGCC |
| H90-92 (3´ substrate) sense | GCTAATACGACTCACTATAGGGATGTCGGCTCTTCCTATCATACCGAAG CAGAATTCGGTAAGCG |
| H90-92 (3´ substrate) anti-sense | CAGCTCACGTTCCCTATTAGTGGGTGAACAATCCAACGCTTACCGAAT TCTGCTTCGG |
