## Supplementary material for "Interprotomer Communication and Functional Asymmetry in H/ACA snoRNPs": Table S4

| **ID** | **Name** | **Sequence** | **Length** | **Effective**  **Length** | **TPM** | **Num**  **Reads** |
| --- | --- | --- | --- | --- | --- | --- |
| 1 | TRINITY_DN2645_c4_g1_i1 | GCTCTTCCGGGGCGGCACATGATTTACGTGCATATTCCAATTTGGAATATGTAACTGCCAGAATGTGCAACAGTAAATAGGGAATTTCGCTAACACCATTCTATCATAGAATGGAGTTTGATAGTACTTCTCGAGACATTTGT | 146 | 4.996 | 75469.4 | 789078.8 |
| 2 | TRINITY_DN6630_c0_g1_i1 | GGCCTGCACTAAATGTCCACATTCTCGGATAATATAGCCGCAGAGTGTTACTGCGAAATAAGTGCAACATTAATGTAGTGACATGGCGTTCTGGACTCGCTCTGTGGTGAGGACGGGTGCAGAGTTATTACAACAAATGAAAAAAAAAAAAAG | 156 | 5.473 | 59296.2 | 679176.6 |
| 3 | TRINITY_DN4609_c2_g1_i1 | GCTCCCACAATTTTGCCCCAGAGCTGAACTTTCTGTGTCAAGATCTGGCTTCTCTGATTGTGGATAGTAAATATTGATTTTAGCTATACCTGACTACATCACCCATTGTAGATTCTGGTACGCAAGTCAATAAACAATTGAAAAAAAAAAAAAGA | 158 | 5.577 | 51783.9 | 604481.7 |
| 4 | TRINITY_DN0_c0_g1_i1 | GGTACTGCAGTCCAGCTATATGAAGAAAAACCATATGGTCTTCATATCGTTTTTGTCTGCAGATATCATATGCTACCGCACATAACCTGCCTCTGTATCTCATATACTTTGGCAACATGTGTCGTTGCTACACGCAAAAAAAAAAAAAGA | 153 | 5.322 | 44891.8 | 500003.9 |
| 5 | TRINITY_DN6637_c0_g1_i1 | GGGTATCCTGTTCATTTGGGAGCCGTACGCCTCTATGGTTGCGGTTCTAGAAAAGACAGGTACATTATATTAGTGAGGCGCCCGTGTTGAGGCGGACTTGATAGACGCCATGCAACCCCGTACGTCTCTCACATTTGGT | 142 | 4.824 | 47803.1 | 482663.9 |
| 6 | TRINITY_DN0_c0_g1_i2 | GGTACTGCAGTCCAGCTATATGAAGAAAAACCATATGGTCTTCATATCGTTTTTGTCTGCAGATATCATATGCTGCCGCACTGATCTATCTCTGTACATTGATGTACTTAGGTACCTTGTGTCGTTGCTACAATCGAAAAAAAAAAAAAT | 153 | 5.322 | 41080.2 | 457550.7 |
| 7 | TRINITY_DN241_c0_g1_i1 | GGATCGCAGACAGCAGTGTGAAGAGCTTAGGTCGTTATTTTTGCTCTTGACATGTGCCTCCTCAAAGCTGCGTACATTATCCGGCGACGCCACGAACGCGGCTCTCGATATTTCGTTGGCTATTCGTGGACCCGCGACACGCAAAAAAAAAAAAAGA | 160 | 5.686 | 30476.9 | 362661.6 |
| 8 | TRINITY_DN1_c0_g1_i1 | GTATCCACATCCAGGCAGTGCCTTCAGTTATATTTCTGTCGGTACGTTCAGATGTTTGTGGGATAGTAACACGCAAACTAGCATATACTCTTTGTTCCTATGAACAAGAGAGTCTACAAGGTTTGCGACATTTGTAAATTAAATATAA | 151 | 5.225 | 29384.0 | 321328.0 |
| 9 | TRINITY_DN2690_c0_g1_i1 | ACTGACTAAAGGGCCTAGCATGGTTCCACAGACGGTGCGCTATGTATAGCTACAGTCTTCTAAGAACATGCTATATTAAAATTTGTGTTTCCTATTAAGGTTTCTTCTTTCATAGAGGAATTCTTAGCGAAGCATATATGACATTTGT | 151 | 5.225 | 29123.5 | 318478.9 |
| 10 | TRINITY_DN1036_c2_g1_i1 | TTCCGATCTGGGGGGTCCAATGTTCGTTAGTGATATATCGAAACACAAGGATACATCACTTCCATAACATTGGAAAATCAAAGATTATAACTGCTCCATTTGTCTCTAGATCTCAAAATATCAGAGTCACGGAATTAAGTTAATACGCAAAAAAAAAAAAAGA | 166 | 6.032 | 21694.8 | 273903.1 |
| 11 | TRINITY_DN1038_c0_g1_i1 | GGCACGGACAAGGGGTTCCCTGATAAAATTATTGTTTTTAAGCTAATATAGTTTAAAAACACCAGTATCAGGGTATAGTATATCGGCGCCGAGCAGAAGATTCAGACTCATGTCTGTCTACTTTTACAGTGTGCTGAACAAGTGAAAAAAAAAAAAAG | 161 | 5.741 | 22613.5 | 271713.8 |
| 12 | TRINITY_DN3_c0_g1_i1 | CCCCGCTCTTCCGATCTGGGGCACCCTACCAAAGCCGATGTGTGCACATAAAAGTTGTGCTCATGTGACTCTGGATAGGAACATTATCACACTCTGTGTAAGACATCCAGTTGTTATAGTTGTATGTCCAAGGGTTGAGTGTGACAACTGAAAAAAAAAAAAAG | 164 | 5.913 | 20971.7 | 259530.0 |
| 13 | TRINITY_DN4601_c0_g1_i1 | ACAAATGTAGGAAGTGTACACAAACCCAACCTTCGAAGATGTCGGTTGAGCAAATTCGCTTCCCAAGTGGATCTTGCATGTAGTTGCAGATGAGTTGACTATTAAATAGTCGACTATCTATGCGTCATGCTTACCCCG | 138 | 4.663 | 25403.9 | 247908.4 |
| 14 | TRINITY_DN240_c0_g1_i1 | GGGGGTAATCCAAATTCGTATATGGTAGCTTGTTAAATTTTTGTTAACGAGAAAACGAATGTGGAAAAAGCAAAGTCTTGACTAGGTGAGGTTGACTGATACACAAATTTTCAGTTAAGCTCAATCCGTCAAGACACAAACAAAAAAAAAAAAAGAA | 160 | 5.686 | 18791.1 | 223606.0 |
| 15 | TRINITY_DN6627_c0_g1_i1 | GGGGGCACATTTGTTCAAGAGTGATGATAATAACAAAGCTTTACTCTGTAAAATATGTGCTATAGTAAACACCATATAATATAGAATAAAATCATGTTTTTTTATTTACATTATTGTGGTGAACAGGCAAAAAAAAAAAAAG | 145 | 4.952 | 17170.7 | 177956.0 |
| 16 | TRINITY_DN4607_c1_g1_i1 | GCATTGCAGCCCAGCTATATGAAAGCTGTAAATCACTTTCATATCGTTTTATGCTGCAAATAATATAGTTGACACTGCTTGGTGAATGTCGCGTGGTTTATTTTTCGCATGGCATTTTCTGTGGGGTCTACAAATGAAAAAAAAAAAAAGA | 154 | 5.371 | 15314.4 | 172159.0 |
| 17 | TRINITY_DN2649_c1_g1_i1 | GCGGGGATTCTCCTTAAAAGCTAACTACTGATCTATATATCTTTAGTTATGTCTATGGAGAGATATTAAGTTTCCCAGCCAACCATCCGACTGAACATCCGTTTTAGATTGGACGTAGTCTGAGGAAAACAAACAAAAAAAAAAAAAGA | 149 | 5.131 | 15707.7 | 168687.0 |
| 18 | TRINITY_DN4605_c3_g1_i1 | CCGCACGACGCTCTTCCGATCGGGGTACGCCACGTTCTCTGGGCAAGAGCAGACAATGTCAATGAGCATACTGACTCTAATCGGATGTGGCAATATTAAATCTCGACGTTAATCCCAAATAATCTTAATTTAAGATATTTGATTCAAGTCAGGGAGACATTTGAAAAAAAAAAAAAT | 177 | 6.769 | 10985.0 | 155615.9 |
| 19 | TRINITY_DN8333_c0_g1_i1 | GAAGCTCCATTCATCCTCTGAATTATAGGTCTTTTCTATTATTTAGACAGCGTTATTGGAGCAACATTATACATTAGGATTTTTGGTTGTGATAAAAATTACACCACTATATTTTTGAAACCGGTCCTAATAACACGTAAAAAAAAAAAAAGA | 156 | 5.473 | 12696.5 | 145424.7 |
| 20 | TRINITY_DN2653_c0_g1_i1 | GCATTGTCTGTCCTAATACCAAGGAGCGTAATATTAGTTCCTTGGCTTCGCTTTCAGACAACATCAAAGGTCTTCAAATTTATCCTAGTGTGATTGTTCTAGGTATGAATATTGCACTGGCATTTTGAAGTTACAATGAAAAAAAAAAAAAGA | 156 | 5.473 | 12307.9 | 140974.3 |
| 21 | TRINITY_DN250_c1_g1_i1 | CGGGGTCGTCCCATTGATACTGGGTAAAGTTTCAATGCTATACACCAGCGGCAGACCGGGACAAACAACAAATCTCAACTGGGATTGGTGAACTGTTATTTGTTGACTTTTTACAACGATTGTTGAGATACATTGG | 136 | 4.585 | 14607.4 | 140183.0 |
| 22 | TRINITY_DN2669_c2_g1_i1 | CTTTTTTTTTTTTTTACATGTATCATACATTTATGGGGTTGTATACTACAATAGCTTACAACCAATTCCGGTATGAGGTATGCTATTAGTCCTCTTTGACACCCAGATTCTTACAAGAAGTAACTAGCAACACTGGGACTGCCCCG | 146 | 4.996 | 12666.9 | 132439.8 |
| 23 | TRINITY_DN8315_c0_g1_i1 | CCCAGCACTCACACAGGATCTCATTTGTAAAATATCGTTTGGGGTCCTGTTTGAAGAGGCAGCAGACTCACTGCCCCATCAAATTTGTGCTCACATTAACATATAAACCACAGGGCCAATGGTGATGAGGACCTTGCCATATGGTAGTATAAGGATTATGACATTGAAAAAAAAAAAAGA | 183 | 7.236 | 8546.3 | 129428.0 |
| 24 | TRINITY_DN2_c0_g1_i1 | GACGCACTAACTGAATTATAAAAATAATGTTTTTTGTAAGCATTTGTTTGGATGTAGTGCAACAGTAACTTGTTTGTCGTACTCACGCGGCCGCTGTCGGCCGATGTGGGCAGTAGCATTCATACAATGAAAAAAAAAAAAAGA | 147 | 5.04 | 12131.3 | 127969.4 |
| 25 | TRINITY_DN6624_c1_g1_i1 | CCGGGGATAACTGGGGGCATCCCCATACATTTGCCACGCTCGCCGTAGCCATTCAATGGCATGTTCGGGTGCCGCGTATGGGGTACATTAACGCTGTCAAGTAGAATACCACTGAGCCACATGCCAGTGTGTATGGAACAATACAGCTACATTTGT | 156 | 5.473 | 11172.3 | 127967.3 |
| 26 | TRINITY_DN6703_c1_g1_i1 | CGGGGTGCCCTGAATTCTTTCCAGACTTTGTTCCATACATTGTTTGGCAGTACTCAGGGAAAATTGAAAAGTATAATTCTAACTGAAATGCATTTTCTATATCATTTTGGTTTTAACATACTGACAGGTAAAAAAAAAAAAAG | 143 | 4.866 | 12179.3 | 124041.7 |
| 27 | TRINITY_DN2662_c0_g1_i1 | GTCTCCCTGTTTAAAACTGACGCAGTACAATACTATGATTGTGGACAGTACAATGGCAGGGTAAATTAAAGTTCTAGTAGCGCGGACGTCAACTCTATGTGGGTTGACGCATTCGATATATTAGAATACATTTGAAAAAAAAAAAAAG | 151 | 5.225 | 11091.5 | 121290.6 |
| 28 | TRINITY_DN6629_c1_g1_i1 | CCGATTGTAGCTGTATTGTTCCATACACACTGGCATGATGCGCAGTGGTATTCTACTTGACAGCATTACTGTACCCCCCACTTGGCACCCGAGCCTGCTAAGAGGAAGCAAGACGAGCGTGGCAATTGTGAGGGGAAGCCCCG | 143 | 4.866 | 11716.1 | 119323.6 |
| 29 | TRINITY_DN4604_c0_g1_i1 | CTTCCGATCTGGGGAATCCACATCCAGGCAGCGCCAATAGTACCAACTACTGTCGGTGCGTTCAAGTGTTCGTGGGACATTAACACGCAAATTAGCATATACTCTTTGTACCCATGGGCAAGAGAGTCTACAAGGTTTGCGACAGGTGAAAAAAAAAAAAAT | 165 | 5.972 | 9033.9 | 112916.6 |
| 30 | TRINITY_DN5_c0_g1_i1 | ATTTTTTTTTTTTTCAACTGTCCGTGGCATGGGTAATGCGCGTGTGAGTACAGACTCACCGCGCGACCTTATGTCACGCTCGCTGTGCGCCGCGATATGACTGACGCGAGGTATGGACTACCTCGCACAGCTTCGAACCGGCGACCCCG | 149 | 5.131 | 10271.7 | 110308.8 |
| 31 | TRINITY_DN6621_c1_g1_i1 | GAAGCACTAGGACAATTCAAAAGTAATGCTTACTCGTATAGTATTATTTACTTGTTTAGTGCAACACTAAATGTGTTTGTCGTACTCACGTGACCGCTGTCGGTCGATGTGGGCAGTAGCATTCATACAACTGAAAAAAAAAAAAAAAG | 152 | 5.273 | 9909.3 | 109358.1 |
| 32 | TRINITY_DN243_c2_g1_i1 | GGCCTTGCACACGATTAGAGCCGTGTACGCGTACACAGCGCTGTGCGGTGTCGAAGTTGTGCTACAGTAAAGTGCCAATCAACCTCGGACCTCGCTCATTGCGGGGCCCGTTCACTGGTGGCGCGACATGCAAAAAAAAAAAAAG | 148 | 5.085 | 9282.6 | 98796.0 |
| 33 | TRINITY_DN4627_c1_g1_i1 | ATTTTTTTTTTTTTCAATTGTCAGAATATTAAGTAATAATACTATCCGTATTTCGAGAACAGTATTAATTTTAATTGAAATTCATTAATTCTTTGTTAATTTAGGAAGCTTTAACTCGGAACTGGTCAAAAAACTGAGAACAGTTCTGTCTCCCATTATCTTCCATGCCCCG | 172 | 6.416 | 7115.6 | 95553.4 |
| 34 | TRINITY_DN4_c0_g1_i1 | GGTGCCTCCGAATTAAATCCCACGTCTACGCCTGTCGTAGCTAGCCGTGGTAGGCCTCGGAGGGACATGAAGATGTCGGTATGACGCGACAGCCTGCACCCACATGAGCATGTTGTGTTATTAATGTCTGCACAAACGAAAAAAAAAAAAAT | 155 | 5.422 | 8178.8 | 92804.0 |
| 35 | TRINITY_DN6622_c0_g1_i1 | TATTTTTTTTTTTTTCGTTTGTCGTATCAGCATTGCACCCGTTCTGACAATACTGAGTCATTCCGGAACGCCCGCCGACACAGTTGGTGTCGCACTCACATTCGCAGTAACTGTGTGCGATATACACACCGCGCACAGTGGACATTTAGTGCATACCCCAGATCGGAAGAGCGGGG | 176 | 6.695 | 6582.5 | 92243.1 |
| 36 | TRINITY_DN2664_c1_g1_i1 | GTGTCCCACTCCCAACGCAGTGTCTTCAGTTAAATATAACTGTGTGTCACATTATAACTTAAGTGGGTATAATAAAAGCTTCACTAAAACACATGCAGGTTCCTGTGAATTTGTGTGCACCTAAGTATTGCGACAATTGAAAAAAAAAAAAAG | 156 | 5.473 | 7911.6 | 90619.0 |
| 37 | TRINITY_DN2658_c0_g1_i1 | CGGGGTACGTCCAACCAAATTACTGTTCATGAATTTACACTTGGACAGTGTCGAAATTGGACAAAAGTAATGCAAAATTATTACGTTTTAAAATATTTTTGTTATTTTTATAAGTATTAAAATTGCAACATTGGT | 135 | 4.547 | 8784.6 | 83607.0 |
| 38 | TRINITY_DN6_c0_g1_i4 | GAACCTGTGAAGTAATATGTCACTGGTAACACCTATGTGATATCAGTATCTGAGCAGGAATATCAATATTTCGTATTTACTCTCGTATGTTTTATGATATATGAGATCGCAAGCGGAATTACACTGGAAAAAAAAAAAAAAG | 145 | 4.952 | 8066.7 | 83602.5 |
| 39 | TRINITY_DN6623_c0_g1_i1 | GTTAGCACTAGCCATTCACAAAGTGCGCATTTCGATGATGCGTCACTTTTTTAAGCGGTGCTAACAGCAACTGCGTGTCGTACTCACCCGACTGATGTCGGCCGAGGTGGCCAGTAGCACTCAAACATGGAAAAAAAAAAAAAT | 147 | 5.04 | 7888.6 | 83214.0 |
| 40 | TRINITY_DN2695_c2_g1_i1 | CCCCCCCCCTGCAGGGGGGACCTGAGGCCATAGAGTCGCGAACTTGGTTACTATACCATGTTTGTGCTGCGGTTACTCAGACATCAAATACCAGACTTTATAAAGCGTCTCATTTTACCCTTGTGTTTTGAGATCTTCGTATAAAGCCATAATTGAAAAAAAAAAAAAG | 169 | 6.219 | 6390.3 | 83180.3 |
| 41 | TRINITY_DN2647_c2_g1_i1 | GAGCCGAGCCCATAGAGTCGCGAACTCGGTCCCATACCGTGTTCGTGCTGAGGGTACTCGGACAGCATATACCAGGCCCGACAATGCGTCTCATTTCACTATCTGCGCTATGAGATCACCGTCCAGGGGCACAGTTGAAAAAAAAAAAAAT | 154 | 5.371 | 7372.2 | 82875.7 |
| 42 | TRINITY_DN4731_c0_g1_i1 | GTGTCCCTGTTATCTTTCAATGCACACTAACACAAGGTTGTGAATTGAACAAGCACAGGGTAAATAAAATGTAATGTTTTTGAGCCGATTTGTGATTCAATCGCACTTCTGTTTGAACCATTACAATATTTGC | 136 | 4.585 | 7524.5 | 72211.0 |
| 43 | TRINITY_DN1037_c1_g1_i1 | ACAAATGTATTTCTTATGTGTCGAAACGTATATTCACAAGAAATATACGTTCGCGACATAAGAAAGTTAATTTCCCCTGCCGTAGTGACCCTCTTTAGTATTTAGAACTTAAGAGGCAAGAAAACAGGGACCCCCCG | 137 | 4.624 | 7412.6 | 71732.9 |
| 44 | TRINITY_DN6646_c1_g1_i1 | GCTTGGATTTGATACAAGTTATAATCTACTCTTAATTATAACTTCACTTGCGATCCATACATTAACTTGTCCTTATGAAGACTTTCTGTTATTAAGCTTTTACAGATAGACAGATACGAGGGCTACATTTGTTT | 137 | 4.624 | 7369.3 | 71313.7 |
| 45 | TRINITY_DN6735_c0_g1_i1 | GATCCTACTCTGATTTAAGATGTATTCACTCTCTAGGATACTTCTTTGTATAATGTAGGTATAGTATTATGAAACCAGATCATGATTAATGTTTTCACTATTACATTTTTCCTCATTGATGGTTTCTGACATTTGC | 139 | 4.702 | 6991.1 | 68802.0 |
| 46 | TRINITY_DN2685_c1_g1_i1 | GAGTCCTGTAAAGATTTTTTCAATTAGTAACACTATGTGTTATTGGTTATCTGAGCAGGTATATCAAAACTTTCGTATTTACTCTCGTATATTTTGTAATGTATGAGATCGCAAGCGAAATTACATTTGT | 133 | 4.473 | 7085.9 | 66340.0 |
| 47 | TRINITY_DN9_c0_g1_i1 | CTCTGCGAGTTATACACGGACATATCATTATAATATTTATTTGATGTGAATCTTTTCACTCGCTACATAATCCACCGATTTGCGCATCAACAACATTGAAGCTTTTGTTGTATTGTGTTTAATATTTGGTGACAGTTGAAAAAAAAAAAAAGA | 156 | 5.473 | 5271.3 | 60377.0 |
| 48 | TRINITY_DN1035_c1_g1_i1 | CTTTTTTTTTTTTTACATGTCTTGAGAGGTACTATCAAACTCCTTTCTATTAACAGAAATGGCGTTAGCGAAAATCTCAATTTATTATAGCACAATCTGGCAGTAACACATTTTATAATAAAATGCGCACGTAAATTATGTGCTACCCCG | 150 | 5.178 | 5496.0 | 59558.0 |
| 49 | TRINITY_DN4600_c0_g1_i1 | GCATTGGATGCAGTACTAGCCGGTTGTTGAATGTTAACAGCTTGGTATATAAGTATCCATACATTAAAAGTCTACTGCCGAAATACAGTTGTTAAGATCTTAAACTGCGTTTGCGATGGAGACTCACAGTTGC | 136 | 4.585 | 6185.3 | 59359.0 |
| 50 | TRINITY_DN6651_c0_g1_i1 | GTAAGTCCCAGTGTTGCCAGTTACTTCAAATAAGCATCTGGGTGTCAAAGTGGACTAAAAGTAAACTTCATGCCGGAATTGGCTGTAAGCTATTGTAGTCTGCAGCCCCATAAACGTATGAGATATTTGC | 133 | 4.473 | 5770.1 | 54021.4 |
| 51 | TRINITY_DN8369_c0_g1_i1 | GCGCATACATTAATAACACAACATGCTCATTTGGGTGCAGGCTGTCGCGTCATGCCGACATCATCATGTCCCTCCGAGGCCTACCACGGCCAGCTAGCGCAGGCTTAGACGTGGGATTTAATTCGGAGGCAGCCC | 135 | 4.547 | 5614.2 | 53433.2 |
| 52 | TRINITY_DN4_c0_g1_i2 | GAAGCCTCCGAAAAATATCCCACGTCTACGCCTGCCCTAGCTAGCCGTGGTAGGCCTCGGAGGGACATGAAGATGTCGGTATGACGCGACAGCCTGCACCCACATGAGCATGTTGTGTTATTAATGTCTGCACAAACGAAAAAAAAAAAAAT | 155 | 5.422 | 4689.3 | 53209.2 |
| 53 | TRINITY_DN13_c0_g1_i1 | GAACGAAGACCCCGATAGAAACACGTTTCTATCCGCTTGACAGCCTGCGTACCGTAACGCCTAAGCGTGCCGCTGATAACGGGCATGCTGTTTCAGGAGCCGTACGTGATGCCGCTCGCCGTTTAATCGACGAGGCTATATCATCCGTCGGAGGATCTAAATTTGAAGTCAACCCGAACCCAAATTCTAGCACTGGATTGCGTAACCATTTCCACTTCGCCGTCGGTGATCTGGCGCAAGACTTTCGCAACGATACACCAGCGGAAGACGCCTTCATCGTCGGTGTTGATGTTGATTACTATGTCACCGAGCCAGATGTGATATTAGAGCACATGCGTCCAGTGGTTTTACACACCTTCAACCCGAAGAAAGTGAGCGGCTTTGATGCTGACTCACCATTCACCATCAAGAACAACCTAGTTGAGTACAAGGTGAGTGGAGGTGCTGCATGGGTTCATCCAGTTTGGGATTGGTGCGAAGCCGGTGAGTTCATCGCCAGTCGAGTCCGAACAAGCTGGAAGGAATGGTTTTTACAACTGCCATTGAGGTTGGTTGGTTTAGAAAAGGTCGGTTACCACAAAATCCATCACTGTAGACCCTGGATTGATTGTCCAGATCGAGCACTCGTCTACACGACACCGCTGTATACCGTTTGGCGGTTCAAGTGGATCGATACCGAGCTGCACGTACGTAAACTGAAGCGGATTAAATACCAGGATGAAACTAAACCTGGTTGGAACAGATTGGAGTATGTCACCGACAAGAACGAACTAATGGTTTCAATCGGTAGAGAAGGAGAGCACGCTCAGGTTACCATCGAGAAAGAAGAGTTGGATATGCTCTCGGGACTATCCGCCACACAATCTGTCAACGCTAGGCTTATCGGTATGGGACACAAGGACCCACAGTACACATCTATGATTGTTCAGTATTACACCGGCAAGAAAGTTGTGTCACCAATCAGTCCGACTGTGTATAAACCTACAATGCCACGCGTCCATTGGCCAATAACCAGTGACGCAGACGTACCAGAAGTGAGCGCACGCCAATACACCTTGCCTATCGTGAGTGACTGTATGATGATGCCAATGATTAAACGCTGGGAGACAATGTCTGAATCAATTGAACGAAGGGTGACCTTTGTCGCAAATGATAAGAAACCAAGCGACAGTATCGCGAGGGTCGCTGACACTTTCGTACGGTTGATGAATGGGCCATTCAACAATCTTGAACCATTGTCAATCGAGGAAACGATTGAGCGTCTGAATAAACCGTCCCAACAACTACAACTTAGGGCGGTTTTCGAGATAATCGGAGTTGAGCCGCGTCAATTGATTGAGTCATTCAACAAAAATGAACCAGGAATGAAATCTAGCCGAATGTATGTCACCGACAAGAACGAACATGGTTTCAATCGGTAGAGAAGG | 1430 | 1180 | 21.2 | 52257.8 |
| 54 | TRINITY_DN8323_c4_g1_i1 | AGAAAAATGTAGCATCTCAATGAATATCTCACACATACTATGCGAGATCAACTTCATATCAACGCATACAATATCCCCCGATTTTGGCACCTGACAACTACTTGTGATATAGTATCAAGTGTGGCATAGAAATGGGGTTGCCCCG | 145 | 4.952 | 4804.7 | 49796.0 |
| 55 | TRINITY_DN1042_c0_g1_i1 | ATTTTTTTTTTTTTTGCAAGTGATAGAGAAAAGAAGGATCTATGAAAACCCTTCTTTATGTTAACCACTGTTGATGTATGCCTTCCGAAAACAGTGATAACAAGAGAGTAGTCACACTGACAAGCTGAAGGCAATCCCC | 139 | 4.702 | 5001.5 | 49222.0 |
| 56 | TRINITY_DN8_c1_g1_i1 | GTTGTCGAGGTCCCTGCGCACGTTGTTTGGCGCGTTCTGGCAGCTCATTCCCATTGCTGCTTCCACCGCCGTTTGAATGCGGTCGGGAAGTTCCTGGATTAGCGCGAGTTTGCTTGGCATCGTTAAGCTTGGGTAACGATTCGTCCACGGAGCCCGGCATAACGCCGTCTTCATCAAACGTACATGCAGCGGCGCTCTCCTCGCTGTCATGTGTAATCTTCTCCCAGACATCCTTCATTGCGGCAAAACGCCCAATGAGTGCATCGACCTGATCTTCGTCAATTCCAGTACGCTTGATTAAAACCTGCTTCATCAAATGGGCGTCTTGCGGGTGCTGTGGCCAGGATCCATTACACGTCAACCAGTAGGGTTTCTCCTTATTCCGGCTTCTACGCTGGTTCCTAACTTCCTCTGTGGAGGCAGTTGGCCCGTATAACCGTAGTACCATTTTGCAATAATCCGATATCAACGGAGTAAGCGCATCGGTACAAAGATAACCTTCGACGCGGTCGCAAGCCGCATCAGCTAGTGGTATCGTAGGATCTCTTGTTGTGAGGTGCAGTTTTCGCAGTGTACGCAGTGGGTCTTGTATTGTTGTCGTCGTTGCGAGCGGATCCACAAAAACACGTGAGAGAAAGCAAAGACCTATCTCTGGGTTGTATCGTTCCACTTTGAGCTCCAAGCCATAGCATTTGGCAGCACGGTTGATAGATTTCCATATGATGGCCCGCGACAGACCATCATCACCGCACTTTGGACCGATTAATCGGAACAGATCTTCTGGTTTGGCAGCTGGGTGCTCAAAGGTGAGAGCGGTAAATTCGACACATGCATTATATTGCGTGTTATGTGGCGTGGTTGTTGGACTTCCACTTTTAACACCAACTCCAGGCTCGTATCGGAAACCAAAGCGTTTAGCTTTGGCTGGACAATGGATGATCGTGTCCATAAATGAAATGATCTCGTCTCTGTATTCTAGCTTGAATGCTTGAACCATCGCCTTTTGAGCGATATTTCGCTGCATCCAGCTGGAAACCCGGCCGTCAAGGTTGGAAAAATCCGTTTCTATGACTTCAGCGTCACAGTCGCTGACAAATTCACAAACCCCGTCGGCGATCTCAGTTGGGTTCCTTCCAGGATAATACCAATGTTTGTTATGCTCGGCATGTAAGACGATATCCGAGTACGCTAATGTGTATCTGGAGACCTTCAGGATGAAAAGGATGTCAGGAAAGCCGGATATTATTCGGCTAGATTTCATTCCTGGTTCATTTTTGTTGAATGACTCAATCAATTGACGCGGCTCAACTCCGATTATCTCGAAAACCGCCCTAAGTCCCC | 1341 | 1091 | 20.8 | 47448.2 |
| 57 | TRINITY_DN1033_c0_g1_i1 | ATTTTTTTTTTTTTCAAATGTTAGATAGAGAACGATGTTACGCTTTAGTGTCTTATGAAGCACAACACAGAGATCTTCTTGTTAATATAGGAAGCGGTTCCGTGTTACCACAGATCATTTGAAAACAATCCACAGAGAATTGCCTCAAATTTAACTTCCATACCCC | 166 | 6.032 | 3714.3 | 46894.0 |
| 58 | TRINITY_DN6654_c0_g1_i1 | GATTGCACTAAATGTCCACTGTGCGCGGTGTGTGTATCGCGCACAGTTACTGCGAATGTGAGTGCGACATGAACTGTGTCGGCGGGCGTTCCGGAATGACTTATATAGTCAGAACGGGTGCAATGTTGATACGACAAATGAAAAAAAAAAAAAG | 157 | 5.525 | 4000.4 | 46255.6 |
| 59 | TRINITY_DN4612_c1_g1_i1 | GTAGGCCTTATCGAAGCTACATGTCGTTTCATACAATATGATAGTCAAATAAAGGCCAATATAAAATGTTCCCGACACGGTCGCGCTTTAGATTCAATTATATCTTCAACGTATCACCGAAGGGGCCTACAATTGAAAAAAAAAAAAAG | 152 | 5.273 | 4071.6 | 44934.0 |
| 60 | TRINITY_DN6631_c0_g1_i1 | GCAAATGTTTGAGTGATAGGAAAGTTGCCCGTGCAAGTAGCACGACACCATAGGGTCACTCGATTTAATGTTGCGCTGTTATTTCGTCACTAGCAACGATGAATATTGATAGTGTTCGTGCTGAGCGCAACCCCG | 135 | 4.547 | 4648.6 | 44243.3 |
| 61 | TRINITY_DN4613_c0_g1_i1 | TCAACTGACTAAGTGGGTTTCCTCATACCTTTGGCATGGGTACGTTTCTACATAGCGTAACATGAATGCGAGTATGAGGTAAATTAAGACAGCTACATGCCTTCGGTGTTCACACAATGTTGTCTGAATCCCATCCTCAGATGCTGTAACATGGAAAAAAAAAAAAAG | 171 | 6.349 | 3313.2 | 44029.0 |
| 62 | TRINITY_DN4646_c1_g1_i1 | TCCCGGGCTTTGCACTAGATGAAGACACAGCATGTAATATACAGTTGTGTTCACCAAAATGGTGCTAAATTATACTAGACTCTCAAAAGTGCTTTATGTAAGTCTCCACACTGCATATTGTTCATTAGAGAGTAACAATTGAAAAAAAAAAAAAT | 158 | 5.577 | 3753.2 | 43812.0 |
| 63 | TRINITY_DN8_c0_g1_i1 | ACGTCTGCGTCACTGGTTATTAAAACCTCTGCCCTTTCGGGCTAGAACGGGTGTGGGTGCTCCTAGGAGTCACCTACGGTTTAGCGGGTGGGGGGTCACTTCCGGTTGTTGGAAGGCTGTGGCTTAGCTCCAGCTTTCGGAGGCGTTCTTCGATTGTTGGTTTTGCCTCCTCGGGGGCCCTTCCTTCGAGGTATGCCACCACGCTGGGTTTCTCCAGCAGTGATGTTACCATTCGGCTTACCGTCAGCTTTGCCTTGTTTAGGCAAGCGTGCAGGTTGTCGAGGTCCCTGCGCACGTTCCCC | 302 | 53.12 | 386.4 | 42954.0 |
| 64 | TRINITY_DN6633_c1_g1_i1 | GGATCCCTCTGAGGCAGTTGTTATCTTTAATAAGACTAACATAAAGAATCAGAGGGAAGATTAATTTTGCTTACTCCACCCTTGTGATAGTAAATTGTATCTCAGGGACTAATATAAAGCATACAAGCAAAAAAAAAAAAAG | 145 | 4.952 | 4129.3 | 42796.0 |
| 65 | TRINITY_DN6_c0_g1_i2 | GACCCTGCGAAGTTACTTCTCACTGGTAACACTTATGTGCTATCGGTTATCTGAGCAGGAATATCAATATTTCGTATTTACTCTCATATGTTTTATAATATATGAGATCGCAAGCGAAATTACACTGGAAAAAAAAAAAAAAG | 146 | 4.996 | 3850.0 | 40254.1 |
| 66 | TRINITY_DN8321_c0_g1_i1 | GCAAATGTTATTTCTGTTAGTATTCTGAACAGTAAAATATAGTGCTCAGAAGTTGCAAATTAGAAATATTACTCTTGCATGAATTTACAAAGAGTAGCACAAATGATGTTACCCTTGAACAAGTTCATGCTGCCCC | 136 | 4.585 | 4010.6 | 38489.0 |
| 67 | TRINITY_DN13_c0_g1_i2 | GAACGAAGACCCCGATAGAAACACGTTTCTATCCGCTTGACAGCCTGCGTACCGTAACGCCTAAGCGTGCCGCTGATAACGGGCATGCTGTTTCAGGAGCCGTACGTGATGCCGCTCGCCGTTTAATCGACGAGGCTACCGAGCTGCACGTACGTAAACTGAAGCGGATTAAATACCAGGATGAAACTAAACCTGGTTGGAACAGATTGGAGTATGTCACCGACAAGAACGAACTAATGGTTTCAATCGGTAGAGAAGGAGAGCACGCTCAGGTTACCATCGAGAAAGAAGAGTTGGATATGCTCTCGGGACTATCCGCCACACAATCTGTCAACGCTAGGCTTATCGGTATGGGACACAAGGACCCACAGTACACATCTATGATTGTTCAGTATTACACCGGCAAGAAAGTTGTGTCACCAATCAGTCCGACTGTGTATAAACCTACAATGCCACGCGTCCATTGGCCAATAACCAGTGACGCAGACGTACCAGAAGTGAGCGCACGCCAATACACCTTGCCTATCGTGAGTGACTGTATGATGATGCCAATGATTAAACGCTGGGAGACAATGTCTGAATCAATTGAACGAAGGGTGACCTTTGTCGCAAATGATAAGAAACCAAGCGACAGTATCGCGAGGGTCGCTGACACTTTCGTACGGTTGATGAATGGGCCATTCAACAATCTTGAACCATTGTCAATCGAGGAAACGATTGAGCGTCTGAATAAACCGTCCCAACAACTACAACTTAGGGCGGTTTTCGAGATAATCGGAGTTGAGCCGCGTCAATTGATTGAGTCATTCAACAAAAATGAACCAGGAATGAAATCTAGCCGAATGTATGTCACCGACAAGAACGAACATGGTTTCAATCGGTAGAGAAGG | 893 | 643 | 28.2 | 37898.1 |
| 68 | TRINITY_DN289_c0_g1_i1 | GTGCGCTCCGCACGAGCGGTGCCAGCCCCATGCTGGTCTCGCAAAAGGAGCGCAAAGTAAACTGTGTGCCCCTATGGTGACCTTCAACATGAGGGATCAACTTTCCTAGCACCCAAACATTGG | 126 | 4.229 | 4188.4 | 37074.0 |
| 69 | TRINITY_DN1043_c0_g1_i1 | GCCGACCCCTGGGACATATCGGCCGCGTTTACAAAGTGTGGCCAAATGCCGCGGGGGTGAGATCGACGCGGCACAACCGAACTGCTCGCGTCACTACGCTGTGCAGATCATACGTGTCGCTACATGTTT | 132 | 4.437 | 3534.3 | 32820.0 |
| 70 | TRINITY_DN4692_c0_g1_i1 | GAACAGGGAATTTATTGAATATCTTTAAAATGTAATTTACGACTGCTAAGACAACATTCTATGAGAGAAAACGTCAAGCTGTCGTAAATATTAATGTTGCATGTTCTTAGAAGACTGTGGCTACGTAGCACACCGACTGTGGAACCATGCTCCCCC | 156 | 5.473 | 2707.6 | 31013.0 |
| 71 | TRINITY_DN4616_c2_g1_i1 | GCTGAGCATTGGAACTTAACCCAATATTCATGCAGTGTGGCACTGTTCTAGTTCCATGCTGTGTGGAAATATCTCATGTGGTGACAACTGTGTGGGATTCACTCTGATGCTAACAGTGACGCACATGCATAGTACCACACTGTTCTAAATGTAAACATTGTGGCCTCCCATATGTGCCACATTTTTTT | 191 | 7.948 | 1830.5 | 30449.0 |
| 72 | TRINITY_DN6639_c0_g1_i1 | GAATGCCTTTAAGTAGATCATTTCGATTTTGTATAAATCGTAATGAGTGGACAAGGAGGCTATATTAACGCCCCGAGCAATGATTCCCAAGTGCATTTAATTATGTATTGGCACTTGCAACGGTTTGTGGCGACATTCGAAAAAAAAAAAAAAG | 157 | 5.525 | 2594.4 | 29998.0 |
| 73 | TRINITY_DN11_c0_g1_i1 | GGTTCGAGTATAGAGTCTCTGATACATTTTGTAATGTGTTGGTGCTAAATCCACTCGTAAATTAAATCTCAGCCCCGTTACCTGGGCTAGTATTTAATTACAAGCCATATGGAAAGTGGGTCTTGAGTACAAATGAAAAAAAAAAAAAG | 152 | 5.273 | 2678.0 | 29553.7 |
| 74 | TRINITY_DN4660_c0_g1_i1 | GGATCCCCAATACTCTTCCTAGTTGTACTCTGTAGTCTTCTAGGTACAATAATAAGGGGTATATTATCATTTTCCCTTAGGAAACACTTTAGTTTTCATGCTAACGGTGTGTCGACATGAGGAATACATTGAAAAAAAAAAAAAT | 148 | 5.085 | 2692.1 | 28652.0 |
| 75 | TRINITY_DN4618_c1_g1_i1 | GATTGCACTAAATGTCCACTGTGCGCGGTGTGTGTATCGCGCACAGTTACTGCGAATGTGAGTGCGACATGAACTGTGTCGGCGGGCGTTCCGGAATGACTTATATAGTCAGAACGGGTGCAATGTTGATACGACAAATG | 143 | 4.866 | 2805.4 | 28572.1 |
| 76 | TRINITY_DN4738_c1_g1_i1 | GGTTCTCCTTAAAAGCTAACTTCAGATCGATATATCTGACGTTATGTCTATGGAGATATATAAATTTTCCAAAGACAACAATCTGATTGAATTATTTCACTCTGACATAGACTATGGAATACATTTGG | 131 | 4.401 | 2917.6 | 26874.8 |
| 77 | TRINITY_DN1045_c0_g1_i1 | GACAGCAAGGGCACTCTGTGTTATTCATTCTATATGATAATTACAATTTGTTACCTTGCTAGAACAAATCTATCACATCCTATTCCTGGCAGCTTGCATCTCGCTGTTTGGTCATCTTGATAGAAACATTTGG | 136 | 4.585 | 2682.2 | 25740.1 |
| 78 | TRINITY_DN2668_c0_g1_i1 | GGCATCCCGTATATTAGGCCTCATTTCTCTATAATAGAGATTCTGTGTTACGCAATACGGGTATAGTAATTTTTTTTGCGAGTAGGTTTTATTGAGGACGTCGCAAACGATGTTTTTGATTACTCGCTGTGAAATACATTGGG | 146 | 4.996 | 2432.4 | 25432.6 |
| 79 | TRINITY_DN2754_c0_g1_i1 | GAAGCATGGTAATATTAGGAGTGAGACATTCAGTGCTCTCTGTTTAGCATACCATGCTAGATCAGAAATCTCCGAACCAAGGTCCTTCCTTCCTTATGAGGAAATGCACACTGATGTGAGATGTACAAGCAAAAAAAAAAAAAG | 147 | 5.04 | 2362.9 | 24925.0 |
| 80 | TRINITY_DN4743_c2_g1_i1 | GGATGCACATATGAAAAATATTTAAATAAACATATTTCTATATTTGCAAGGAACAAACTGTGCAATAATAAACTGATGGTCAACTAGACACCGGCTCATGCTCCTATGCTAGATTCGGTCGCTTTATACCTACAGTTTATATGTTAATGTACCTGG | 159 | 5.631 | 1979.1 | 23324.0 |
| 81 | TRINITY_DN7_c0_g1_i1 | GTATCCCCCCCATCCCGTTGCTAACCTCAATAGGTTTGACAACTGCGTACAAGGGGGTAAAGTAAAGTAAGCATTGGAAGCGTCGCATAGTTCTTAACGGCTAGCGGCGTGCCGAAAATGTTTATACAATCGG | 136 | 4.585 | 2314.6 | 22213.0 |
| 82 | TRINITY_DN1085_c0_g1_i1 | GTTTGCACAACTACCGCGGATCGCCAATAGTTCTTGTTGGTAGACATTTGATACGTTGTGCTATATTAAATAAAATGTCACAAAAGGATATCTATCTCCTGTGGTAGATATTGTTAGTGGTGTTTTAGACATTTGAACAGTTGTTGTG | 151 | 5.225 | 1921.9 | 21016.8 |
| 83 | TRINITY_DN4606_c0_g1_i1 | GGAGGCATCATCTAATACCACTGGTTGATCATTAATATCATCCTATGGAGCTATTGATGCAAGAGCATATAACTTCAGTGAAAAAACATCCATTCTGTAATGTTTGTTCATGAAAGGGGGTTGACAAATGAAAAAAAAAAAAAT | 147 | 5.04 | 1968.6 | 20766.0 |
| 84 | TRINITY_DN1041_c1_g1_i1 | CTTTGCACTGGGTATCTCCACGGTTGATAATATATCGATCGTGGTCGTTTATACTGTGCTATAATAAATATGCTCACCCATTCGAAAACACAAGCTACTCTCTGTTTTCGTCAGTTACGAGCATTACATTTGG | 136 | 4.585 | 1880.7 | 18049.0 |
| 85 | TRINITY_DN37_c0_g1_i1 | CGCGTTAATCAGCCACAGATAACAACACAAAATGTCACGAGTTGAGTTATTCTGTGGTTAGAGAAGGAGCAAAAGAAACATTTCATACAGCCTAATGTGTCAAACACAAATATTATGGACGGATATTGGTGGTTTAATTGGGTGATTCAATTTTGGGACACTATGCGTTTAAGGGTTAATGAAGAAAACGAAAACGCGACAGAACATCGTCCGTCGTCGCACACGTTGTCTTGATGTGTGTTTGTGTGTTTACATTATCATCGTCCTCGGACGAATCACCTGGCGTAGGGCTGAGTCTCAACAGATCGCAGCACGACGCTGCTCTACCGAGCACAACACCCCGCCAGGAACGGAAGTCGTCTACAGACTATTCCGAGCCCCGACATCGAACTGAGGTAAATTCGGACCTTCGGAGCCGTGATGCACGCGTTAAACGGACAGCATCGATCTCCGCGATCCAAATGGGCTTCGACGTCGCACCTCACGTGGTGAAGCGCGACTAGTAAAGTCACATTGTTTAGAGCCTCCCGACTCTCGGGGCTCCACAGTGAGCATATCCTTGCCGGATTCGGCTAGGCTGGCTTCGGCCTTAGAGGCGTTCAGGCATAATCCCGCGGATGGTAGCTTCGCACCACCGGCCGCTCGGCCGAGTGCATGAACCAAATGTCCGAAACTGCGGTTCCTCTCGTACTGAGCAGTATTACTATCGCAACGACAAGCCATCAGTAGGGTAAAACTAACCTGTCTCACGACGGTCTAAACCCAGCTCACGTTCCCTTTTGATGGGTGAACAATCCAACGCTTGGCGAATTTTGCTTCGCAATGATAGGAAGAGCCGACATCGAAGGATCAAAAAGCAACGTCGCTATGAACGCTTGGCTGCCACAAGCCAGTTATCCCTGTGGTAACTTTTCTGGCACCTCTTGCTAAAAACTCTTTATACTAAAGGATCGATAGGCCGTGCTTTCGCAGTCCCTATGCGTACTGAACATCTGGATCAAGCCAGCTTTTGCCCTTTTGCTCCACGCGAGGTTTCTGTCCTCGCTGAGCTGGCCTTAGGACACCTGCGTTATTCTTTGACAGATGTACCGCCCCAGTCAAACTACCCGCCTGGCAGTGTCCTCGAACCGGATCACGCGGGAGTTTTACGGCGACGAGCGTTGCCGCCACGTCGCCACTCTGCACGCTTGGAACGAAACACCGTGCGCCCGCCGATTAACATCGACCGCGCACCGCTTCCGCCCAACCGAGTAAGTAATGAAACAATGAAAGTAGTGGTTTTTCAGCGACGATCGCGCGAACGATCTCCCACTTATGCTACACCTCTCATGTCTCCTTACAATGCCAGACTAGAGTCAAGCTCAACAGGGTCTTCTTTCCCCGCTGATTCTCCCAAGCCCGTTCCCTTGGCTGTGGTTTCGCTAGATAGTAGATAGGGACAGTGGGAATCTCGTTAATCCATTCATGCGCGTCACTAATTAGATGACGAGGCATTTGGCTACCTTAAGAGAGTCATAGTTACTCCCGCCGTTTACCCGCGCTTGCTTGAATTTCTTCACGTTGACATTCAGAGCACTGGGCAGAAATCACATTGCGTCAACACCCGCGAGGGCCATCGCAATGCTTTGTTTTAATTAGACAGTCGGATTCCCCTTGTCCGTGCCAGTTCTGAGCTGACCGTTGAACGGCGGTCGTACAGAACCGCGCCGATCGCGCACGAGTCGAGACCGACACGGCCTTACGGCTAGGAAGATCCGCGGAAGGCCGGAACGCGGGTCCGGATTCCGCCCTCGCGCCCCGAAAGACGCGAGAGCATCGACCAGGCCCGGCACCGGCCGCATCCGCTTCCCGTCCAAACCCGACACGCCCCGGTCCTCAGAGCCAATCCTTATTCCGAAGTTACGGATCCAATTTGCCGACTTCCCTTACCTACATTATTCTATCGACTAGAGGCTCTTCACCTTGGAGACCTGCTGCGGATATGGGTACGAACCGGCGCGACATCTCCACGTACATCCCTCACCTGAATTTTCAAGGTCCGCAGAGAGTATCCGGACACCGCCGCAAATGCGGTGCTCTTCGCGTTCCGAACCATATCTCCCTTCTATAGGATTCCATGGAACTCGAACGCTCAGGCAGAAAAGAAAACTCTTCCCGGACCTCTCGGCGGCGTCTTCAGGCCACTTTGGGTTACCCCGTCGAACACTCGCTGTAAAAACGAGGGAACGATTATTGAAACGGTTCCGCTGCCGGGTTCCGGAATAGGAACCGGATTCCCTTTCGCTCAAAGGGCGTTATTTATCATTATATACCATAATAAAATGATACATTATTGAAAAAAACACGCCACATCGACATAAGATTTCTCCTTGAGCTTAGGATCGACTGACTCGCGAGCAACTACTGTTCACGCGAAACCCTTCTCCACGTCAGTCCTCCAGGGCCTCGCTGGAGTATTTGCTACTACCACCAAGATCTGCACCGACGGAGGCTCCAAGCGGGCTCACGCCCAGACCCTTCTGCGCTCTCCGCCGCGCACGTCCTACTCGTTACGGCTTAATGACGTCACAAAGAACGTCGCACATGCCCGTAACGGTAGTGTATAGGCAAAACGCTTCAGCGCCATCCATTTTCAGGGCTGGTTGCTTCGGCAGGTGAGTCGTTGCACACTCCTTAGCGGATTCCGACTTCCATGGCCACCGTCCTGCTGTCATGAGCGACCAACGCCTTTCATGGTGTCCCATGAGCGTTTTTTAGGCGCCTTAACACTACGTTTGGTTCATCCCACAGCGCCAGTTCTGCTTACCAAAATTGGCCCACTTGGCACCGTCATCAGATCTCCGGCTTCATCGTTCGAGTAAGCCGGAGTACTCACCCATTTAAAGTTTGAGAATAGGTTGAGGTCGTTTCGGCCCCAATGCCTCTAATCATTCGCTTTACCGGATGAGACTGTTCAAATCGACGCCAGCTATCCTGAGGGAAACTTCGGACGGAACCAGCTACTAGATGGTTCGATTAGTCTTTCGCCCCTATACCCAGTTCCGACGATCGATTTGCACGTCAGAATCGCTACGGTCCTCCATCAGGGTTTCCCCTGACTTCGACCTGACCAGGCATAGTTCACCATCTTTCGGGTCCCAGCATTTATGCTCAGAGCGCGCCTGCATTCACGGATTGGAAACGAGACGCCTCGGGAGTGCGAGAGATCGACCTAGATCGACGCTCCATCCTCCCTGAGCACGCGGCTAGCGCGCGCCTTCACTTTCGTTGCGCCTTTCAGTTTTATTATCTCAATGACTCGCATACATGCTAGACTCCTTGGTCCGTGTTGCAAGACGGGTCCTGCGAGTGCCCGAAACTGAATCATCGCAGACAGAGACGCGCACAGTCCGTGACAACACGGCTGCGACGACAGACGCGCCGCACCTACGTCCGCACTTTGGCGTAGACGTTGGATACGACGTTGACTTGCGTCGGGCCGGACGCGTCTAAAACGCGTGCGCGATTCGTCAGAACTACCGTCCGACGGCCGGTCGGCCACCGTCGCGGTCCCACTACCCCCCCCG | 3566 | 3316 | 2.6 | 17825.4 |
| 86 | TRINITY_DN6743_c0_g1_i1 | GATCCCGTATTTTAGGCCTCATATTTCTGTCAAATAGAGATGTTGGGTTACGCAATACGGGTACAGTAAATTTCATTCTAAGTAGGTTTTATTGAGAACGTCCCGATTGGATGTTTTTGATTACTTATTGTGAAATACATTTGCAAAAAAAAAAAAGA | 161 | 5.741 | 1475.3 | 17727.0 |
| 87 | TRINITY_DN8411_c0_g1_i1 | CATCGCACGCTCCTTAGATCCGTTTCATTTTCGATGTGAAACGCTTGAGTCATAAGTGTGCAAGAGCATAATACCTCCATACTGAGACTGATATGTGATTTTCTTAAGATGTTCACTTGTCTCGGAATGGGGAACATTTGT | 144 | 4.909 | 1697.1 | 17435.0 |
| 88 | TRINITY_DN1073_c0_g1_i1 | GCGTGTCGCGGGTCCACGAATAGCCAACGAAATATCGATAGCCGCGTTCGTGGCGTCGCCGGATAATGTACGCAGCTTTGAGGAGGCAAATGTCAAGAGCAAAAATAACGACCTAAGCTCTTCACACTGCTGTCTGCGATGCC | 143 | 4.866 | 1660.2 | 16908.6 |
| 89 | TRINITY_DN2_c1_g1_i1 | GTAGCACCGCCTGCTTAACAAATAACGTATTGATACCGTGCGTTATTATCAGTCAGTGCAACAGTAACGTGTTTGTCGTACTCACGCGGCCGCGGTCGGCCGATGTGGGCAGTAGCAT | 121 | 4.068 | 1779.2 | 15150.3 |
| 90 | TRINITY_DN2712_c0_g1_i1 | AGAGTCTCGTTCGTTACCGGAATTAACCATACAAATCGCTCCACCAACTAAGAACGGCCATGCACCACCACCCACCGAATCAAGAAAGAGCTGTTAATCTGTCAATCCTTCCGGTGTCCGGGCCTGGTGAGATTTCCCGTGTTGGGTCAAATTAAGCCGCAGGCTCCACTCCTGGTGGTGCCCTTCCGTCAATTCCTTTAAGTTTCAGCTTTGCAACCATACTCCCCCCGGAGTCCAAAATCTTTGGTTTCCCGGAAGCTGCCCGCCGAGCCATTGTAGTAACGTCGGCGGATCGCTAGATGACATATTTACGGTTAGAACTAGGGCGGTATCTAATCGCCTTCGAACCTCTAACTTTCGTTCTTGATTGATGAAAACACCTTTGGCAAATGCTTTCGCTGATGTTCGTCTTGCGACGATCCAAGAATTTCACCTCTAACGTCGCAATACGAATGCCCCCAGTTATCCCTATTAATCATTACCTCGGAGTTCTGAAAACCAACAAAATAGAACCGAGATCATATTCTATTATTCCATGCACGAAATATTCAAGCAGCATTTTGAGCCCGCTTTGAGCACTCTAATTTGTTCAAAGTAAAATTGTCGGCCCACCTCGACACTCACCGAAGAGCACCGCGATAGGATTTTGATATTGAACCGGCGTATTACCGCCGGCTCACCGACGATATGCTCCGCAGACGTGTCAGTATCACCGCGGATGCGGTGCACCGACAGCGCGGCGCACAAATGCAACTACGAGCTTTTTAACCGCAACAATTTTAGTATACGCTATTGGAGCTGGAATTACCGCGGCTGCTGGCACCAGACTTGCCCTCCAATTGTTCCTCGTTAAAATATTTAAAGTGTACTCATTCCGATTACGAGGCCTCGTAAGAGTCCCGTATCGTTATTTTTCGTCACTACCTCCCCGTGCCGGGAGTGGGTAATTTGCGCGCCTGCTGCCTTCCTTGGATGTGGTAGCCGTTTCTCAGGCTCCCTCTCCGGAATCGAACCCTGATTCCCCGTTACCCGTGACAACCATGGTAGTCGCAGAAACTACCATCGAAAGTTGATAAGGCAGACATTTGAAAGATGCGTCGCCGGTACTGGACCATGCGATCGGCAAAAGTTATCCAGATTCATCAAAATTAACGACTTCAGACACATGGCCATCCGTCGATTGGTTTTGATCTAATAAAAGCACTCATCCCATCACTGGTCAGAGTTCTGATTGCATGTATTAGCTCTAGAATTACCACAGTTATCCAAGTAACTGAGTAAGATCTAAGGAACCAAAACTGATATATTGAGCCATTCGCGGTATCGCCTTAATACGGCTTGCACTGAGACATGCATGGCTTAATCTTTGAGACAAGCATATAACTACTGGCAGGATCAACCAGGGAGCTTCATTCATATGAAAGACGTTCTCGTTCGTTTCGATTTTCGATTGCGGTGGTACACACGTATACCGTCCTCTCTCTATATCGATACTTACGAGAAATCCTCATTCCCGCGCGTCGACGGATTAACGCAGACGCTTGTACGGGCGAGCTCGTATAATAATTGAGACGAATATTGTTTTCGACGACGACGACGACGACTACGACGACGCCTATAAGGGTACAATATAATGAAAAAACCCCGATGGTGAACGTTCGTATCGAGACACGATTTCATTCACTTTAATAATAATCTCGGTTAACATACCC | 1712 | 1462 | 4.7 | 14386.8 |
| 91 | TRINITY_DN6641_c0_g1_i1 | GGACCCTGTTCTTACTGTTTCTAAGATACCCAAATCTTAGCAATTATATGCGCCAGGGGAGAACAACTGGTATTGAAGTAATCCTGTGGGTTATTTGTCCAAAGGCATAATTTGATACCATACATTTGC | 132 | 4.437 | 1538.9 | 14291.0 |
| 92 | TRINITY_DN8317_c0_g1_i1 | GTATGGAAGTTACAACTAGTTGGGGACAATATCATGTTGTTTTGTCACCGAATCACCTTCCTATATCATCGAGAAGATATCTGAGCGGCGTCTCTTAAGTACTAAGACGTTGTTTCGTTCCCCTTCACACATTTGAAAAAAAAAAGAAA | 152 | 5.273 | 1283.3 | 14162.0 |
| 93 | TRINITY_DN8575_c0_g1_i1 | CTCTGTCTCGTTAGTTAATAAGAATAGGTCTTTTGAGTTATTCTAGAACACGGGACAGATATTGACGTGGTCAAAACTGCCGGCGAAGGCGCCGACGCCGCCACTCCGGTTATATAGACACACTACAATTGAAAAAAAAAGAAAA | 148 | 5.085 | 1272.6 | 13545.0 |
| 94 | TRINITY_DN1103_c0_g1_i1 | GTATGCACACTTTGATCTATCGCAGTTGTAACAGCAGCTGCGGTAATAACAGTTGTGCTAGATAACTGCGGCTGTATCCACACATGTGCGCGACTTGCGTAACTTGTACTAATTTTGTCGCTACATTCGAAAAAAAAAAAAAG | 146 | 4.996 | 1284.5 | 13430.0 |
| 95 | TRINITY_DN4639_c0_g1_i1 | CACCCCACGTTTTTCCTGAGCGGACTAGCATGCACACGATAGCTCCGCCTTTCCGAGACGTGGCAGATTAACAGTTCTTTGTTTTGGCATCGTCACCCGACATACACCTTGCCTGAATAGGGGAACTAACACGCAAAAAAAAAAAAAT | 151 | 5.225 | 1171.7 | 12813.0 |
| 96 | TRINITY_DN10_c0_g1_i1 | GCAAGCACCGTCGATCAGTGCCCACGCCCATGCGTGGCCCACGTTTGCAGCGGTGCGAGAGTAGTCGCGGTACCAGCTAGTTGCGTGCACGTTGCGCGCCGCTTTCCTAGAAACCGCGGGACATTTCT | 131 | 4.401 | 1317.6 | 12136.5 |
| 97 | TRINITY_DN6751_c0_g1_i1 | CTCTTCCGATCTGGGACTCCAATGTTCGTTAGTGGTGTGCACATATTTAGATAAGCTCATCACTTCCATAGCATTGGGAGATAAAATGCGGCAATTGCTCCGTTTCCTTCTACTTAATAGAAAGTCACGGATCTCAGTTCGCAGACATTGG | 154 | 5.371 | 1060.9 | 11926.0 |
| 98 | TRINITY_DN1137_c0_g1_i1 | GATCTCCATTTCCGTTCTATAAGTTGACTGAAGACATTATCACCTTATTTTTAAGATGGAGTACATTATTGGATATTTGTGGTGAATTAAATCGATGCGTCACTTCTAGCATCGATTTATGAACACTGTGTCCGACATCTAAAAAAAAAAAAGA | 157 | 5.525 | 1015.6 | 11743.5 |
| 99 | TRINITY_DN24_c0_g1_i1 | GTATTAAATTAATTCCGTGACTCTGATATTTTGAGATATAGAGACAAATGGAGCAGTTATAATCTTTGATTTTCCAATGTTATGGAAGTGATGTATCCTTGTGTTTCGATATATCACTAACGAACATTGGACCCCC | 136 | 4.585 | 1141.5 | 10954.7 |
| 100 | TRINITY_DN8374_c0_g1_i1 | GTATTGCATCGTCTTGCTCCATGCACTTAACACACAGGGTGCTTGGGATAACTAGATGCATAAACTATGGCCAATGCGTCGCTCTTTGTAATCACTAATGGTTACCGAGATGTACCGCAATGGCACAGTCGAAAAAAAAAAAAAAG | 149 | 5.131 | 988.1 | 10611.0 |
| 101 | TRINITY_DN6669_c0_g1_i1 | GTTGGACTAAGAAGGGTTATTTAATTCTTAATTATTAAATAATCTTAATTATCTAAAGTCCAAAATTATGTGATACAAGATAGGCTTGCTTATTCCGTAATTTGTAAGGTTGAGAAAGTATCAAACATTTG | 134 | 4.51 | 1074.2 | 10140.0 |
