## Supplementary material for "Interprotomer Communication and Functional Asymmetry in H/ACA snoRNPs": Table S5

**Table S5: Cryo-EM data collection, model refinement and validation statistics**

**Data statistics**

| Sample | | Cross-linked snoRNA | Cross-linked  snoRNA | Cross-linked snoRNA | Cross-linked snoRNA |
| --- | --- | --- | --- | --- | --- |
| Microscope | | Titan Krios | Titan Krios | Titan Krios | Titan Krios |
| Voltage (kV) | | 300 | 300 | 300 | 300 |
| Electron exposure (e^–^/Å^2^) | | 60 | 60 | 60 | 60 |
| Pixel size (Å) | | 0.414 | 0.414 | 0.414 | 0.414 |
| Symmetry imposed | | C1 | C1 | C1 | C1 |
| Initial particle projections | | 7,128,691 | 7,128,691 | 7,128,691 | 7,128,691 |
| Final particle projections | | 760,070 | 733,250 | 134,682 | 74,734 |
| Map resolution (Å) | | 2.92 | 2.95 | 3.14 | 3.23 |
| FSC threshold | | 0.143 | 0.143 | 0.143 | 0.143 |
| Map resolution range (Å) | | 2.5 - 7.0 | 2.5 - 7.0 | 2.5 - 7.0 | 2.5 - 7.0 |
| EMD code | | EMD-XXXXX | EMD-XXXX | EMD-XXXX | EMD-XXXXX |
| Structure model  PDB code | | Class I  XXXX | Class II  XXXX | Class III  XXXX | Class IV  XXXX |
| Model resolution (Å)  FSC threshold | | 2.92  0.143 | 2.95  0.143 | 3.14  0.143 | 3.23  0.143 |
| Model composition  Non-hydrogen atoms  Protein residues  RNA base pairs  Ligands  Zn ions | | 12320  1272  101  0  0 | 11840  1225  96  0  0 | 10251  1080  76  0  0 | 11363  1167  97  0  0 |
| *B* factors (Å^2^)  Protein  RNA  Ligands and Zn | | 67.63  109.20  --- | 81.25  121.24  --- | 86.91  124.20  --- | 64.89  106.84  --- |
| R.m.s. deviations  Bond lengths (Å)  Bond angles (°) | | 0.004 (0)  0.921 (0) | 0.004 (0)  0.620 (0) | 0.004 (0)  0.708 (0) | 0.007 (0)  0.674 (5) |
| Validation  MolProbity score  Clashscore  Rotamer outliers (%) | | 0.99  0.71  0.18 | 1.54  4.88  1.02 | 1.36  3.37  0.53 | 1.99  7.19  1.86 |
| Ramachandran plot  Favored (%)  Allowed (%)  Disallowed (%) | | 96.17  3.82  0.00 | 95.93  4.07  0.00 | 96.42  3.58  0.00 | 94.26  5.74  0.00 |
| CC (mask) | | 0.78 | 0.67 | 0.63 | 0.75 |
